# Glutamate/GABA Resource Asymmetry Enables Bidirectional Activity Correction

**DOI:** 10.64898/2026.11.16.767034

**Authors:** Xiaoyu Zhang, Qianhui Chen, Yifei Zhang, Mingyi Huang, Xiangkai Lian, Yuguo Yu, Dewen Hu, Xin Lu

**Author notes:** Corresponding authors: Dewen Hu and Xin Lu.

## Abstract

Glutamate/GABA cotransmission allows one neuron to excite and inhibit the same target. How the two release branches translate a shared spike train into net synaptic drive, and how this drive shapes neural circuit activity, remain unclear. We introduce BRIDGE, a model with separate glutamate and GABA resources that deplete and recover under shared input. The model reproduced glutamatergic depression during repeated stimulation in co-transmission recordings. Late GABAergic responses showed weaker frequency dependence. This asymmetry can generate negative differential gain in the model, allowing faster presynaptic firing to weaken net drive while output remains excitatory. In simulated circuits, cotransmitting neurons shifted their output toward excitation when activity fell and toward inhibition when it rose, improving recovery after perturbations across all circuit sizes tested. Compared with baseline, a simplified spiking implementation yielded higher mean accuracy in most classification comparisons and lower error in selected prediction tasks. Thus, resource asymmetry links firing history to bidirectional correction of circuit activity.

## 1 Introduction

Synaptic communication shapes information processing and collective activity in neural networks. Understanding how synapses maintain responsiveness to inputs while limiting excessive recurrent amplification is therefore a central problem in circuit function. Dale’s principle describes the consistency of a neuron’s transmitter phenotype across its axonal outputs [1, 2]. Many network models represent this organization by assigning each neuron exclusively excitatory or inhibitory outputs, whose strengths vary with recent activity through short-term synaptic dynamics [3, 4]. In this framework, the regulation of network activity depends on interactions between distinct excitatory and inhibitory populations [5–7].

However, multiple experimental studies have shown that glutamate/GABA cotransmission enables a single presynaptic neuron to provide both excitatory and inhibitory signals [8, 9]. Evidence from identified pathways and multiple brain regions further indicates that cotransmission is widely distributed [10, 11]. Moreover, changes in the relative transmitter contributions alter postsynaptic firing and circuit activity [12–14].

Specifically, Root et al. mapped neurons co-expressing glutamatergic and GABAergic markers and identified concentrated populations in the ventral tegmental area, entopeduncular nucleus, and supramammillary nucleus [9]. Their ultrastructural analysis showed that individual terminals from these populations formed both asymmetric and symmetric synaptic contacts. Within lateral habenula terminals, vesicular transporter labeling distinguished glutamatergic and GABAergic vesicle populations. These transmitter-specific pools were associated with different synaptic specializations, placing separate excitatory and inhibitory release sites within a single presynaptic output.

Hirai et al. examined the release properties of these two components at supramammillary inputs to dentate granule cells [15]. Minimal-stimulation recordings resolved separate gluta-matergic and GABAergic quantal responses, while measurements of calcium sensitivity, presynaptic modulation, and calcium channel–vesicle coupling revealed distinct release properties. Both components showed short-term depression, but increasing stimulation frequency produced stronger depression of glutamatergic responses than of late GABAergic responses. This difference changed the relative excitatory and inhibitory contributions during repeated input, enabling frequency-dependent filtering of supramammillary signals.

The relative contributions of the two transmitters also depend on the postsynaptic target and can undergo selective long-term modification. Ajibola et al. found that supramammillary inputs produced different excitation–inhibition ratios across dentate granule cells and interneuron classes, with excitation predominating in dendrite-targeting interneurons and inhibition in soma-targeting interneurons and granule cells [16]. These inputs sharpened granule-cell spike timing, enhanced granule-cell spiking in response to cortical input, and promoted long-term potentiation at cortical–granule-cell synapses. Tabuchi et al. showed that granule-cell depolarization selectively potentiated the glutamatergic component of supramammillary cotransmission, increasing its relative contribution and promoting granule-cell firing [17]. This potentiation was expressed postsynaptically through AMPA receptor recruitment and did not require NMDA receptor activation. Hirai et al. subsequently identified a Hebbian form of selective glutamatergic potentiation induced by pairing supramammillary input with granule-cell spikes [18]. This input-specific plasticity required NMDA receptors and CaMKII and strengthened the ability of supramammillary and perforant-path inputs to jointly drive granule-cell firing.

Changes in transmitter contribution can also alter activity in other cotransmitting circuits. Shabel et al. showed that GABA co-released from basal ganglia inputs limits lateral habenula firing [12]. The relative GABA contribution decreased in a depression model and increased after antidepressant treatment, linking the composition of a common presynaptic output to habenular excitability. Meye et al. found that cocaine withdrawal reduced presynaptic vesicular GABA transporter (VGAT) expression at pallidal inputs to the habenula, weakening GABAer-gic cotransmission and increasing habenular activity [13]. Selective VGAT overexpression at these terminals restored GABAergic transmission and alleviated withdrawal-associated behavioral effects. Pelkey et al. demonstrated glutamate/GABA cotransmission from hippocampal CCK-positive, VGluT3-expressing interneurons to principal cells [14]. Compromising GABA synthesis shifted their normally inhibition-dominated output toward excitation and promoted network hyperexcitability.

The timing of transmitter action also shapes the postsynaptic response. In the medial habenula projection to the interpeduncular nucleus, Ren et al. found that brief stimulation evoked fast glutamatergic currents, whereas stimulus trains recruited a slower current mediated mainly by nicotinic acetylcholine receptors [19]. In cortical circuits, Galarreta and Hestrin showed that differential short-term depression changes the relative efficacy of excitatory and inhibitory synapses during sustained activity [20]. For glutamate/GABA cotransmission, Krubitski et al. modeled how differences in the timing and amplitudes of AMPA and GABA_*A*_ components interact with dendritic filtering and ionic currents [21]. Their simulations produced diverse excitatory, inhibitory, and transient summation patterns, demonstrating how a common co-transmitting input can shape postsynaptic temporal processing. Together, these experimental and computational findings highlight the role of cotransmission in regulating neuronal activity, while changes in transmitter contributions implicate it in disease-related circuit dysfunction [12, 13]. However, how distinct release dynamics translate a shared presynaptic activity history into changes in net synaptic gain, and how this gain regulation influences recurrent network activity, remains incompletely understood.

To address this question, we introduce BRIDGE (Branch-Resolved Independent Dynamics for Gain and E/I control), a dual-resource model that connects distinct transmitter dynamics to activity-dependent synaptic gain [22, 23]. Each presynaptic spike recruits glutamatergic and GABAergic branches with separate resources that are depleted by release and replenished between spikes. Different depletion and recovery dynamics allow the two contributions to evolve differently under a shared spike train, making net synaptic drive depend on the preceding activity history. We use this formulation to determine when increasing presynaptic activity reduces net drive, deriving the conditions under which the steady-state input–output slope crosses from positive to negative. We then connect this negative differential gain to input sensitivity and local stability in recurrent networks. Fits to cotransmission recordings constrain the excitatory and inhibitory dynamics, finite-site simulations test the reliability of the resulting output, and matched recurrent networks evaluate bidirectional activity correction and recovery after perturbations. We further assess the computational utility of the same resource mechanism in trainable spiking networks. Together, these analyses connect release asymmetry and firing history to synaptic gain regulation and recurrent activity control.

## 2 Results

### 2.1 Asymmetric transmitter resources generate negative differential gain

Figure 1 connects the biological motivation to the proposed mechanism and its evaluation across scales. Glu/GABA cotransmission and published electrophysiological recordings motivate separate transmitter resources (panels a,b). Their asymmetric dynamics translate shared presynaptic activity into distinct branch responses (panel c), producing negative differential gain in the illustrative regime (panel d). The co-release motif is embedded in recurrent connectivity (panel e) to study activity correction, and trainable SNNs evaluate classification and temporal prediction (panel f).

**Figure 1.**
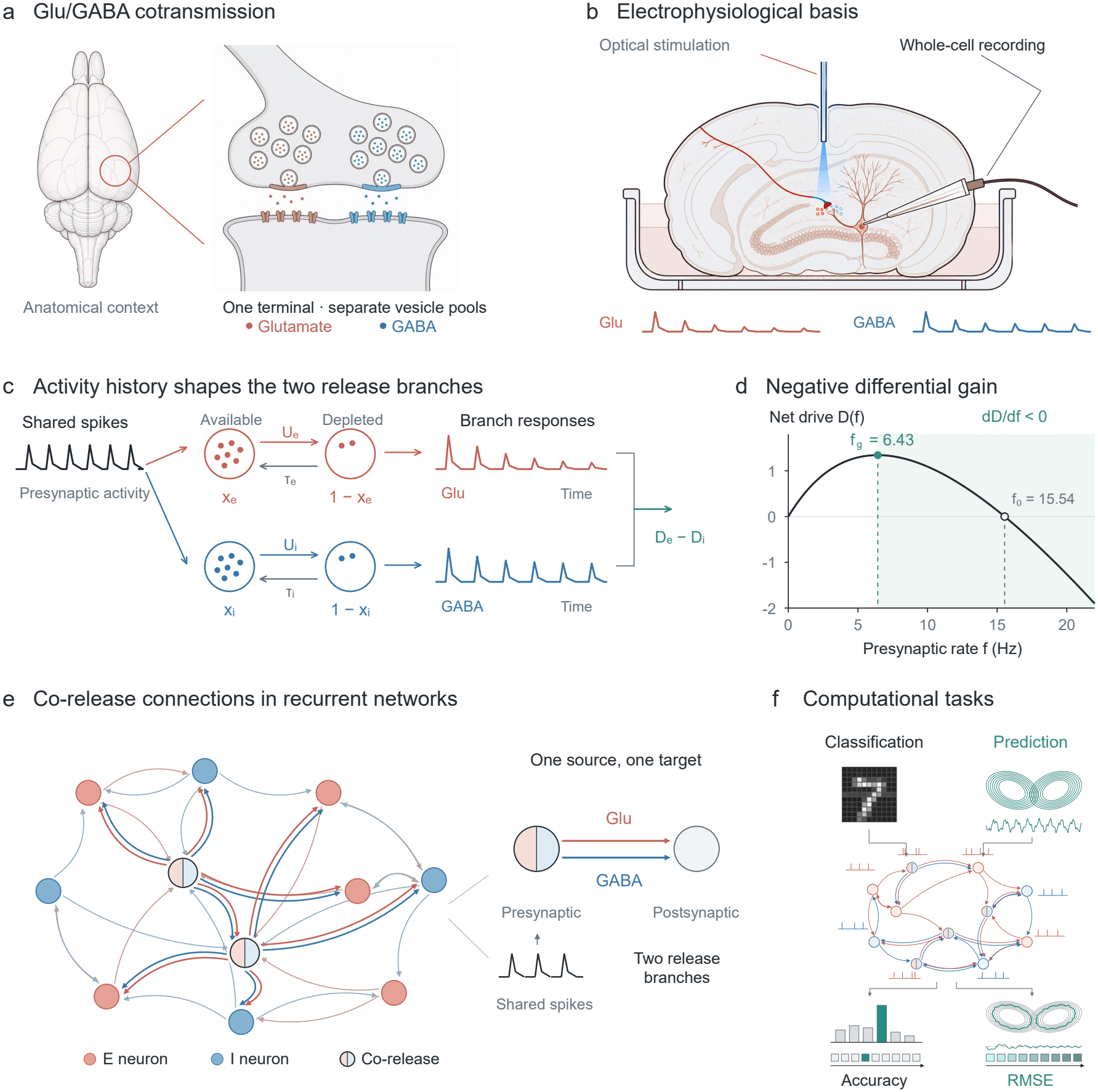
Biological motivation, resource-dependent gain, and multiscale evaluation of BRIDGE.(a) Schematic anatomical context and a common presynaptic terminal with separate Glu and GABA vesicle populations. (b) Optical stimulation and whole-cell recording illustrate the published electrophysiological basis [15]; data and fitted parameters appear in Fig. 2. (c) Shared spikes drive separate available and depleted resources, with branch-specific depletion *U*_*b*_ and recovery time *τ*_*b*_. Distinct branch responses combine into net drive. Response trains in b and c are schematic. (d) Illustrative drive *D*(*f* ) = *A*_e_*f/*(1 + *α*_e_*f* ) − *A*_i_*f/*(1 + *α*_i_*f* ), using *A*_e_ = 1.005, *A*_i_ = 0.5, *α*_e_ = 0.065 s and *α*_i_ = 0. Shading marks negative differential gain above *f*_*g*_ = 6.43 Hz, preceding the polarity crossing at *f*_0_ = 15.54 Hz. (e) Recurrent network with interspersed E neurons (red), I neurons (blue), and cotransmitting neurons (split-color). Each cotransmitting output is drawn as paired Glu and GABA branches from the same presynaptic neuron to the same postsynaptic target. Red and blue arrows denote excitatory and inhibitory outputs, respectively, with arrowheads indicating transmission direction; the inset enlarges this shared-source, shared-target organization. Self-connections are excluded, as in the simulated networks. (f) Classification and temporal prediction evaluate trainable co-release SNNs, using accuracy and RMSE, respectively. Task inputs and readouts are schematic.

Hirai et al. reported distinct release properties for Glu and GABA, with stronger frequency-dependent weakening of the Glu branch [15]. We first asked what functional consequence follows from this biological architecture. For the illustrative parameter set in Fig. 1d, BRIDGE’s signed drive reaches a maximum at *f*_*g*_ = 6.43 Hz. Above this point, increasing presynaptic rate reduces net drive even though the response remains Glu-dominant. Net polarity reverses only later, at *f*_0_ = 15.54 Hz. This example illustrates how asymmetric branch dynamics can generate negative differential gain while net drive remains excitatory.

We next isolated whether this behavior follows from cotransmission itself or from its resource architecture. When the same branch amplitudes were scaled by one common gain, net drive saturated but its marginal gain remained positive at every finite rate. The closed-form thresholds matched numerical roots across 800 parameter draws, and the continuous-rate classification agreed with the exact periodic model in 97.0% of 672 conditions. Thus, gain turnover arises when independently scaling transmitter branches exhibit sufficient depression asymmetry.

Biological inputs are neither perfectly periodic nor noise free, so we next tested whether the signature survives temporal and finite-site variability. Poisson and bursty renewal trains shifted the location of *f*_*g*_ but retained the turnover (Supplementary Fig. S2c). Finite-site simulations detected the transition in all 150 separate-resource repetitions at *N* = 10–100, whereas false detections in the monotone null declined from 18% at *N* = 10 to zero at *N* = 100 (Supplementary Fig. S2d). The mechanism is therefore robust to input statistics, although threshold estimation becomes less reliable at small site counts.

Release architecture also changes fluctuations without changing the mean. At fixed marginal release probabilities, feasible positive Glu–GABA event correlation reduced net-current standard deviation by 15.2% through cancellation of opposing events. These calculations convert the reported synaptic organization into a testable prediction: sustained high-frequency activation should progressively lose its excitatory impact. We next asked whether the measured synaptic trains and postsynaptic output show this behavior.

### 2.2 Biological recordings support the predicted frequency dependence of cotransmission

We first quantified the transmitter dynamics that constrain the model. The SuM→DG EPSC trains showed rapid initial depression followed by a frequency-dependent residual response (Fig. 2a). The resource model reproduced this train profile (*R*^2^ = 0.919), with fitted *U* = 0.857, *τ*_x_ = 1.09 s, and *b* = 8.76%. The downstream analysis propagated both observationally equivalent interpretations of the residual term: a measurement floor and a non-depressing component.

**Figure 2.**
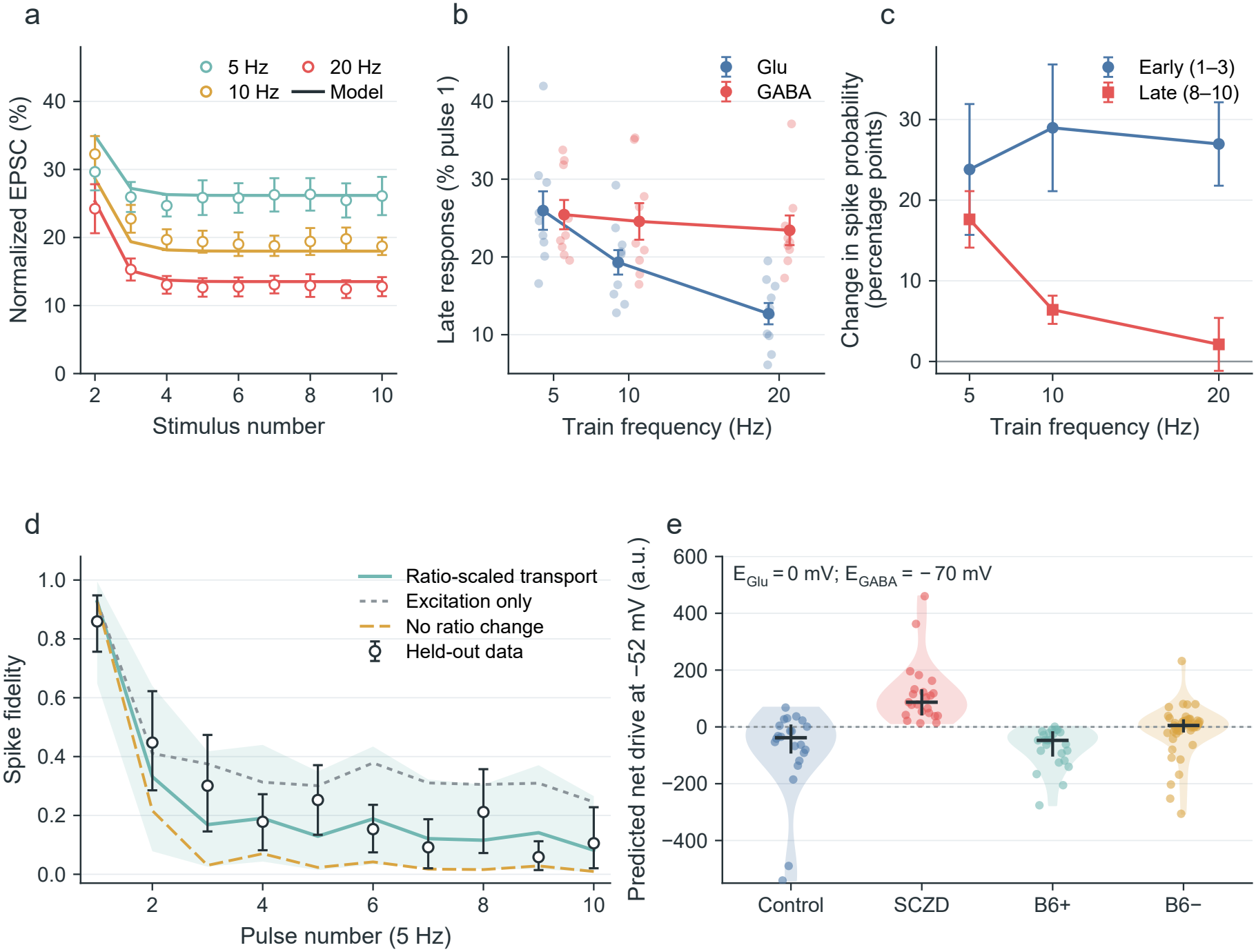
Biological recordings support differential branch dynamics and frequency-dependent output filtering. (a) Hirai et al. EPSC trains [15]; solid curves show the fitted model and symbols are reported means ± SEM. (b) Cell-level late-train Glu and GABA responses at 2.5 mM Ca^2+^; small points are cells and large symbols are means ± SEM. The inferential comparison is the HC3-robust branch-by-log_2_-frequency interaction. (c) Measured increase in spike probability produced by adding intact SuM co-release to the control current drive (Light−control), shown for early (pulses 1–3) and late (pulses 8–10) train epochs; symbols are paired-cell means ± SEM. (d) Fully held-out cocaine+ACSF spike-fidelity train from Meye et al. [13]. The prediction transports a saline-derived inhibitory pulse template using the independently measured cocaine picrotoxin train and GABA_*A*_R/AMPAR ratio; shading is the cell-resampling 95% interval. Excitation-only and unchanged-ratio predictions were fixed comparators. (e) Parameter-free signed-drive prediction from Pelkey et al. conductances [14] at the stated voltage and reversal potentials. Violins and dots show cells; bars show medians and interquartile ranges. Panels a–c connect synaptic dynamics to measured output filtering; panels d and e evaluate transport of branch-strength and output-direction predictions.

We next quantified the branch asymmetry required for gain turnover. As train frequency increased from 5 to 20 Hz, the late Glu response fell from 26.0% to 12.7% of pulse 1, whereas the corresponding GABA response changed from 25.4% to 23.4% (Fig. 2b). The cell-level GABA slope was −1.01 percentage points per frequency octave (*p* = 0.470), whereas the Glu slope was −6.63 points per octave. Their difference was −5.62 points per octave (95% CI −9.56 to −1.68; branch-by-frequency interaction, *p* = 0.0052). The measured trains therefore establish stronger frequency-dependent weakening of the Glu branch while allowing a shallow GABA slope. This divergence predicts strong spike recruitment at train onset followed by reduced recruitment during sustained high-frequency stimulation. The early Light−control spike-probability increment remained 23.8–29.0 percentage points across frequencies, whereas the late increment fell from 17.6 points at 5 Hz to 2.1 points at 20 Hz (Fig. 2c; *p* = 0.0037 for the late 5-versus-20-Hz contrast). Blocking GABA_*A*_ receptors restored a positive high-frequency contrast, linking the reduced late output to sustained inhibition.

Uncertainty in the functional branch weights and fitted dynamics was propagated to the model-derived gain-turnover distributions (Supplementary Fig. S6).

Finally, we evaluated whether branch-selective output predictions transport beyond the discovery dataset. In Meye et al. [13], a saline-derived inhibitory template and independently measured cocaine inputs predicted the fully held-out cocaine+ACSF spike-fidelity train with lower error than excitation-only or fixed-ratio controls (RMSE 0.083 versus 0.149 and 0.162; Fig. 2d). In Pelkey et al. [14], measured conductances predicted net inhibition in control and B6-sufficient cells, excitation after direct GAD inhibition, and a shift toward excitation after B6 deficiency without parameter fitting (Fig. 2e). The SuM measurements anchor the frequency-dependent dynamics, while the Meye and Pelkey analyses extend branch-strength and output-direction predictions across conditions and pathways.

### 2.3 Independent release converges to deterministic dynamics as site number increases

For independent release, the large-*N* analysis predicts that fluctuations in current normalized per site diminish as the number of sites increases. Biological terminals contain finitely many release sites. We therefore varied the number of sites per branch from 10 to 50 to assess how closely stochastic output follows the deterministic prediction. At 10 Hz, the stochastic mean followed the deterministic response at every tested site count, while trial-to-trial dispersion narrowed markedly from *N* = 10 to *N* = 50 (Fig. 3a). In the same representative biological parameter draw, the fraction of release events with net drive opposite in sign to the deterministic prediction decreased from 3.4% at *N* = 10 to zero observed events at *N* = 30 and 50 (Fig. 3b).

**Figure 3.**
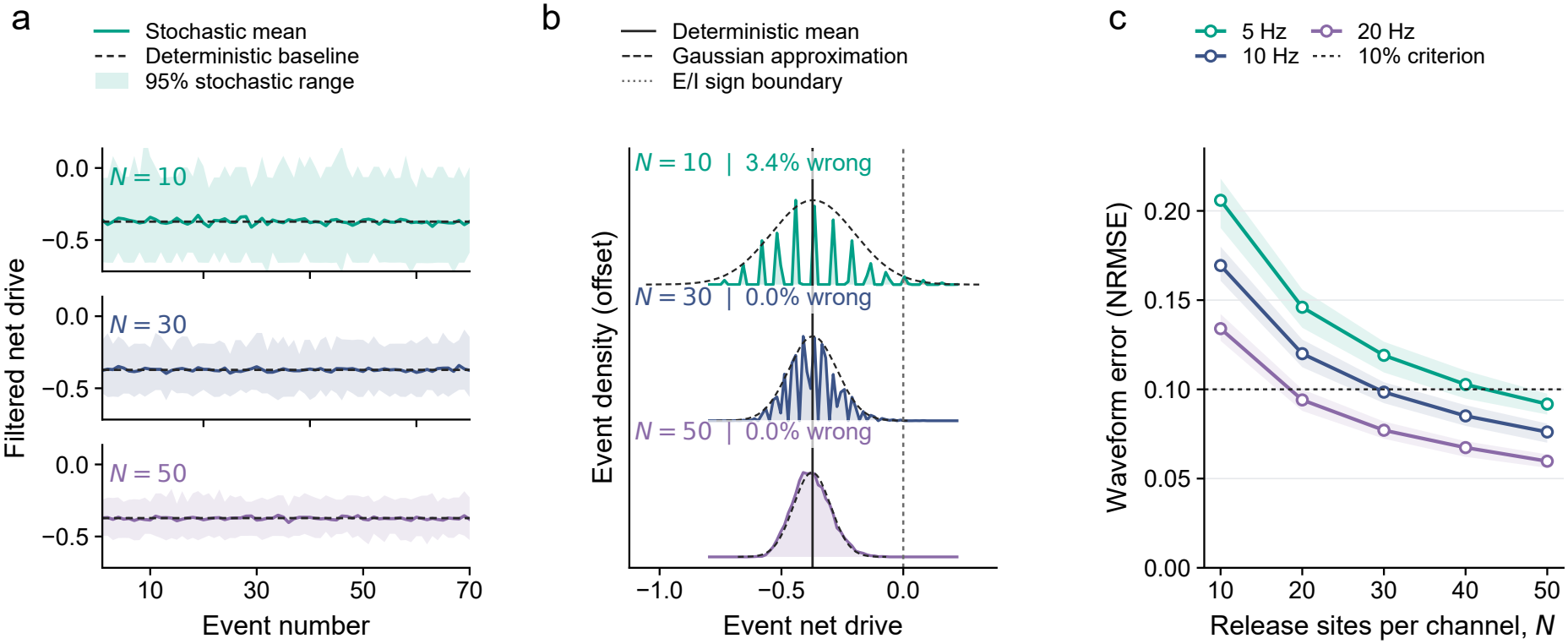
Finite-site release concentrates around the deterministic synaptic output. (a,b) Direct comparisons for the biological bootstrap draw nearest the joint parameter median (*U* = 0.8572, *τ*_x_ = 1107.5 ms) at 10 Hz, using 80 fixed-seed realizations for each displayed *N* . (a) Mean filtered stochastic response (colored line), central 95% trial range (shading), and deterministic response (black dashed line). (b) Empirical event-net-drive distributions (colored profiles) with Gaussian approximations (black dashed curves). The solid vertical line is the deterministic mean, the dotted vertical line is the E/I sign boundary at zero, and the warm-colored tail marks events with the opposite sign from the deterministic prediction; annotations give their observed fractions. (c) Median filtered stochastic–deterministic NRMSE across 100 biological bootstrap parameter sets at each of three frequencies; shading denotes the central 95% interval and the dashed line marks the 10% error criterion. The combined waveform-error and polarity criterion is shown in Supplementary Fig. S7.

Across 100 biological bootstrap parameter sets at 5, 10, and 20 Hz, filtered stochastic– deterministic waveform error decreased with site count at every frequency (Fig. 3c). These results quantify how finite-site fluctuations narrow around the predicted synaptic output. The combined waveform-error and polarity criterion is reported in Supplementary Fig. S7. We next examined how the resulting activity-dependent synaptic drive shapes recurrent responses to perturbations.

### 2.4 Negative differential gain supports bidirectional activity correction

The preceding analyses characterized the synaptic input–output relation generated by asymmetric resource dynamics and quantified its reliability under stochastic release. We next asked whether negative differential gain can provide corrective feedback in recurrent networks, opposing both increases and decreases in population activity. To connect the synaptic mechanism to this circuit function, we combined analysis of recurrent input sensitivity with perturbation tests in reduced and explicit spiking networks.

We first connected differential synaptic gain to recurrent input sensitivity. Across matched working rates, the BRIDGE steady-state slope became negative above *f*_*g*_, and the normalized equilibrium susceptibility fell below one on the stable branch (Fig. 4a). At the matched 5-Hz working point, *χ/q* was 0.436 for BRIDGE and 1.439 for common gain. These values follow Eq. 16 and agree with the response calculated directly from the Jacobian. The full Jacobian also located the stable working range, incorporating the finite recovery time of the resources.

**Figure 4.**
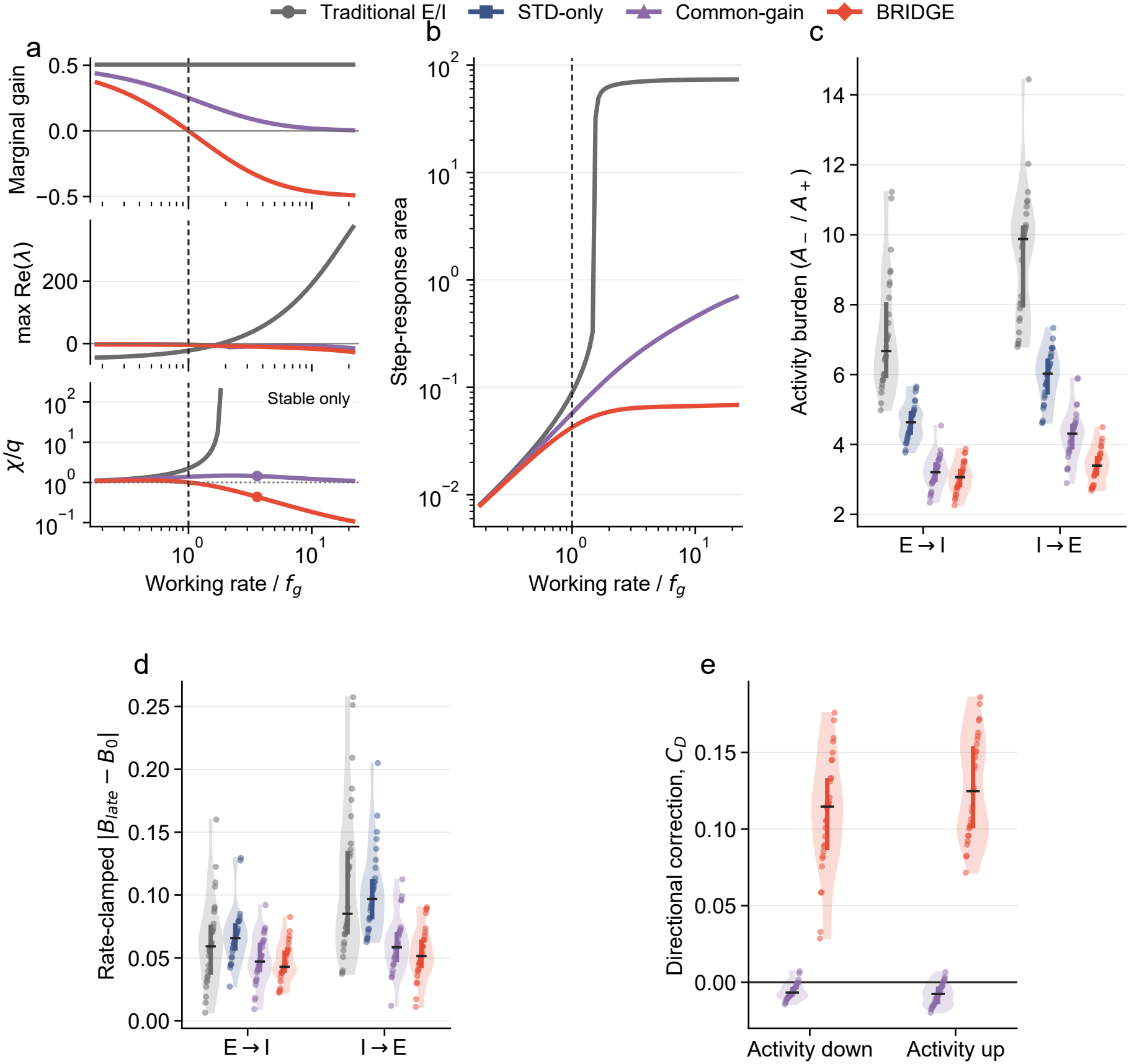
Differential synaptic gain links input sensitivity to bidirectional activity correction.(a) Differential synaptic gain (top), dominant Jacobian eigenvalue (middle), and normalized equilibrium susceptibility *χ/q* on stable branches (bottom) across matched working rates in the reduced recurrent model. The reference at one indicates susceptibility equal to the local neuronal slope; filled markers identify the 5-Hz working point. Rates are normalized by the BRIDGE turnover frequency *f*_*g*_; the vertical dashed line marks *f* = *f*_*g*_. (b) Absolute nonlinear step-response area for symmetric positive and negative input steps around the same operating points. (c) Direction-specific activity burden after permanent 4% E-to-I or I-to-E output reassignment in four baseline-matched explicit networks. (d) Late recurrent E/I error after a rate-only controller returned each trial toward its own pre-conversion firing rate. Population firing rate was the controller’s sole feedback signal. In c,d, violins show *n* = 30 seed-matched networks, dots are seeds, thick vertical bars are interquartile ranges, and black ticks are medians. (e) Directional corelease correction *C*_d_ after pure negative or positive intrinsic-drive perturbations. Positive values denote a co-release shift opposing the imposed activity change; common gain and BRIDGE were branchwise matched at 5 Hz (*n* = 30).

Nonlinear perturbations showed the corresponding reduction in activity excursions. Below *f*_*g*_, median step-response areas were similar across models (0.019–0.022 spikes per neuron); above *f*_*g*_, they were 0.066 for BRIDGE, 0.262 for common gain, and 29.4 for the static model (Fig. 4b). The steady-state slope and the resource-dependent feedback dynamics thus connect the synaptic gain transition to the network response.

We then tested the direction of the local corrective output by changing intrinsic drive. BRIDGE shifted its co-release contribution toward excitation after activity was reduced and toward inhibition after activity was increased (median *C*_d_ = 0.115 and 0.125). The corresponding common-gain values were −0.0067 and −0.0077 (both paired Holm-adjusted *P <* 10^−8^; Fig. 4e). These opposite responses to negative and positive drive steps demonstrate the bidirectional correction predicted around the negative-gain operating regime.

We next imposed a permanent mismatch by converting 4% of recurrent outputs from E to I or from I to E. All four architectures were matched to the same 5-Hz firing rate and *B*_0_ = −0.52 working point, and the common-gain control was additionally matched branch by branch to BRIDGE at 5 Hz. BRIDGE produced the smallest median direction-specific activity burden in both perturbations (3.07 spikes per neuron after E-to-I and 3.40 after I-to-E; Fig. 4c). Relative to common gain, the paired median reductions were 0.16 and 0.91 spikes per neuron, respectively (Holm-adjusted *P* = 0.043 and *P <* 10^−8^). Thus, allowing the two transmitter branches to change relative to one another improved activity recovery beyond cotransmission with a single matched gain state.

Because a smaller rate excursion can itself reduce recurrent-current error, we then clamped firing rate with a controller that observed rate alone. All architecture–direction combinations met the prespecified late-rate error criterion. Under this clamp, BRIDGE retained lower late E/I error than traditional E/I and excitatory STD in both directions (Holm-adjusted *P* = 0.026 and 6.4 × 10^−5^ for E-to-I; 1.9 × 10^−6^ and 1.1 × 10^−8^ for I-to-E; Fig. 4d). BRIDGE and common gain produced similar global E/I error in both directions (both adjusted *P* = 0.617). Rate control therefore preserves the E/I benefit relative to conventional and excitatory-STD architectures, while the common-gain comparison locates the distinctive BRIDGE effect in directional local correction and activity burden.

The mechanism also persists under biological parameter uncertainty: 14,999 of 15,000 joint Hirai bootstrap draws retained negative marginal gain at 5 Hz, and 14,997 retained bidirectional correction, and the activity benefit persisted when recurrent networks were rematched to the inhibition-dominated DG range reported in vivo [24] (Supplementary Figs. S2b and S5). BRIDGE and common gain yielded similar global E/I error at the most inhibition-dominated working point.

### 2.5 Compensation persists across perturbation strengths and network sizes

The time courses reveal how the corrective response develops after permanent output reassignment (Fig. 5a,b). Following E-to-I conversion, the co-release contribution shifts toward excitation; following I-to-E conversion, it shifts toward inhibition. These opposing shifts accompany smaller deviations of population rate and recurrent balance than in the traditional and excitatory-STD controls. The response therefore tracks the direction of the imposed mismatch throughout the post-conversion interval.

**Figure 5.**
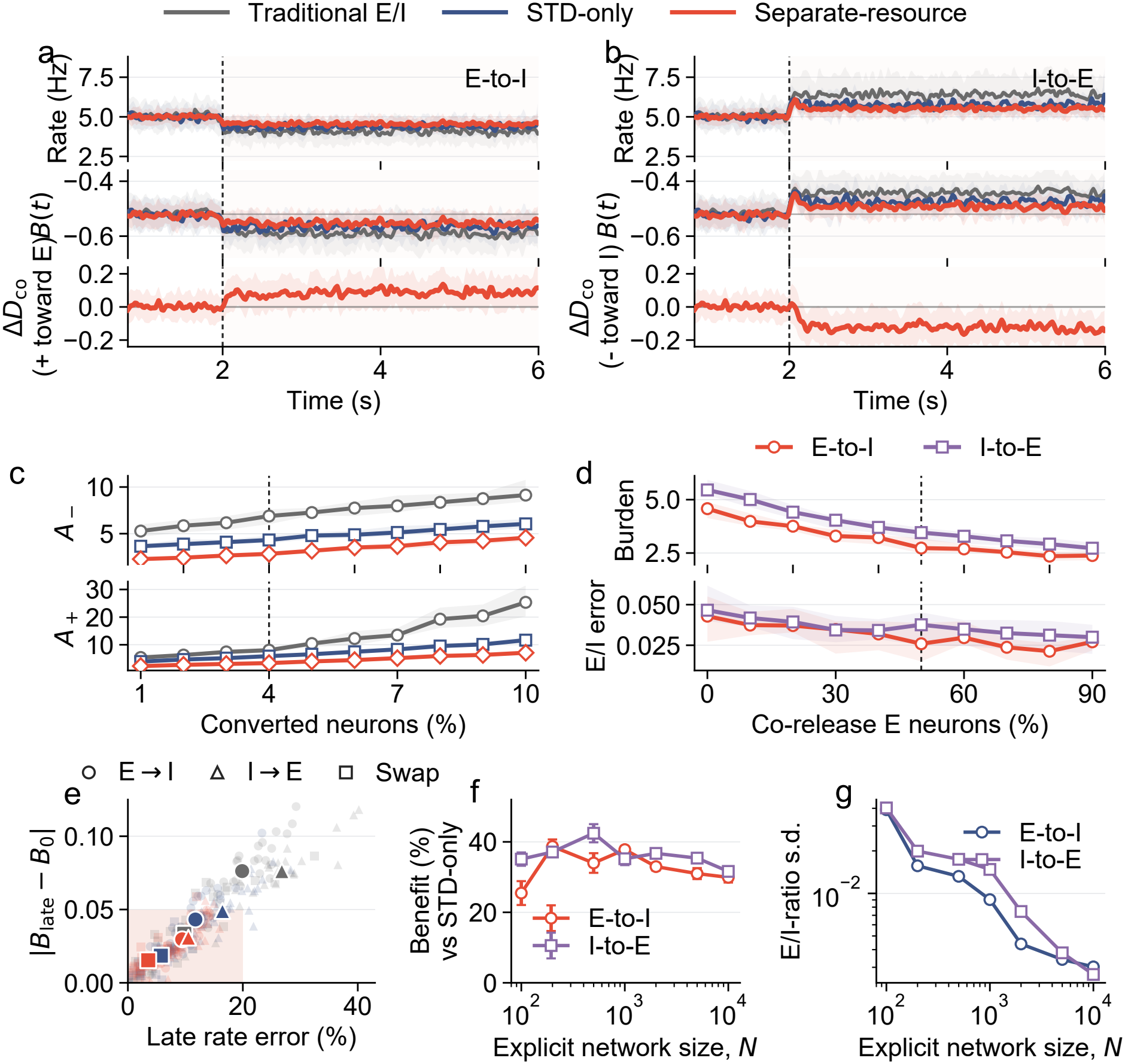
Compensation persists across perturbation strengths, co-release fractions, and network sizes. (a,b) Population rate, recurrent balance *B*(*t*), and BRIDGE co-release drive after E-to-I and I-to-E conversion, respectively, at the frozen 4% challenge. Lines are means and shading is ±1 s.d. across *n* = 30 seed-matched networks. (c) Direction-specific activity burden across every integer conversion fraction from 1% to 10%; lines are medians and shading is the interquartile range. The dashed line marks the frozen primary point. (d) Activity burden (top) and late E/I error (bottom) as co-release prevalence varies from 0% to 90% of E neurons at fixed 4% conversion (*n* = 30 new networks per point); every prevalence was independently baseline matched. (e) Joint late-rate and E/I error for E-to-I, I-to-E, and balanced-swap perturbations; pale symbols are seeds and large symbols are condition means. (f) Mean ± SEM seed-paired burden reduction relative to STD-only across explicit fixed-outdegree networks. (g) Standard deviation of the late E/I ratio across the same 12 seed-matched BRIDGE networks at each size and conversion direction, calculated from the late balance as *E/I* = (1 + *B*)*/*(1 −*B*) over the simulated range *N* = 100–10^4^.

The benefit extends beyond the primary 4% perturbation. Across conversion fractions from 1% to 10%, BRIDGE yields lower direction-specific activity burden than traditional E/I and excitatory STD in both directions (Fig. 5c). Increasing the fraction of co-release neurons from 0% to 90% of the excitatory population also reduces activity burden after each prevalence is independently matched to the same baseline (panel d). Late E/I error follows a less monotonic dependence on co-release prevalence. Joint rate and E/I errors place these effects in a common outcome space for both conversion directions and balanced swaps (panel e). The prevalence sweep characterizes the architectural dose response over the tested range.

Compensation also persists as network size increases. BRIDGE reduces activity burden by 25.5–42.5% relative to seed-matched excitatory-STD controls across explicit sparse networks from *N* = 100 to 10^4^ (Fig. 5f). Across the 12 BRIDGE network realizations at each size, the late E/I ratio varied less between realizations in larger circuits (panel g). Over the seven sizes tested, log–log fits gave approximate power laws for the between-network standard deviation, *N* ^−0.54^ after E-to-I conversion and *N* ^−0.57^ after I-to-E conversion.

### 2.6 Cotransmission improves classification and selected time-series predictions

The preceding analyses show how separate transmitter dynamics support bidirectional correction of recurrent activity. Resource depletion and recovery also retain information about preceding spikes, so synaptic output depends on both current input and recent activity. This temporal dependence raises a further question: can the same synaptic computation improve the processing of task-relevant information in a learned network? We therefore incorporated the co-release motif into trainable SNN units and evaluated classification and time-series prediction across feedforward, recurrent, and gated backbones.

At *α* = 0.5, co-release augmentation increased mean classification accuracy in 31 of 32 dataset–backbone combinations and tied in the remaining comparison (Table 1). The median gain was 3.06 percentage points, with the largest improvements on event-rich auditory and gesture datasets. The motif is therefore compatible with feedforward, recurrent, and gated backbones, although its benefit varies across tasks.

**Table 1:** Classification accuracy and gains from co-release augmentation. Values are mean ± s.d. accuracy (%). The final column reports BRIDGE minus baseline in percentage points (pp). Each condition contains *n* = 4 seeds for SNN and SRNN and *n* = 5 for LSTM and GRU.

| Dataset | Backbone | Baseline | BRIDGE | Accuracy gain |
|---|---|---|---|---|
|  |  | *α* = 0 | *α* = 0.5 | (pp) |
| P-SMNIST | SNN | 19.31 ± 0.95 | 19.96 ± 0.20 | +0.65 |
|  | SRNN | 21.73 ± 0.26 | 21.92 ± 0.07 | +0.19 |
|  | LSTM | 21.86 ± 0.26 | 23.67 ± 0.73 | +1.81 |
|  | GRU | 22.85 ± 0.41 | 24.47 ± 0.82 | +1.62 |
| S-MNIST | SNN | 34.25 ± 0.85 | 34.25 ± 1.77 | 0.00 |
|  | SRNN | 46.43 ± 1.19 | 49.07 ± 1.74 | +2.64 |
|  | LSTM | 49.31 ± 2.02 | 55.49 ± 0.62 | +6.18 |
|  | GRU | 46.51 ± 2.04 | 51.50 ± 1.58 | +4.99 |
| SHD | SNN | 65.80 ± 3.01 | 74.76 ± 0.90 | +8.96 |
|  | SRNN | 69.05 ± 1.62 | 79.87 ± 2.09 | +10.82 |
|  | LSTM | 72.93 ± 4.63 | 80.11 ± 2.99 | +7.18 |
|  | GRU | 47.43 ± 6.80 | 79.51 ± 1.98 | +32.08 |
| SSC | SNN | 33.23 ± 1.80 | 41.36 ± 0.30 | +8.13 |
|  | SRNN | 44.35 ± 1.27 | 55.55 ± 1.38 | +11.20 |
|  | LSTM | 49.39 ± 4.64 | 60.06 ± 0.70 | +10.67 |
|  | GRU | 49.76 ± 2.42 | 61.17 ± 0.27 | +11.41 |
| MNIST | SNN | 96.37 ± 0.12 | 97.55 ± 0.07 | +1.18 |
|  | SRNN | 96.06 ± 0.37 | 97.51 ± 0.03 | +1.45 |
|  | LSTM | 96.33 ± 0.15 | 97.61 ± 0.11 | +1.28 |
|  | GRU | 96.19 ± 0.10 | 97.41 ± 0.03 | +1.22 |
| Fashion-MNIST | SNN | 85.05 ± 0.36 | 87.66 ± 0.12 | +2.61 |
|  | SRNN | 83.94 ± 0.24 | 87.45 ± 0.14 | +3.51 |
|  | LSTM | 85.17 ± 0.14 | 87.53 ± 0.16 | +2.36 |
|  | GRU | 83.82 ± 0.83 | 87.29 ± 0.20 | +3.47 |
| N-MNIST | SNN | 95.32 ± 0.20 | 96.23 ± 0.23 | +0.91 |
|  | SRNN | 93.92 ± 0.33 | 96.01 ± 0.15 | +2.09 |
|  | LSTM | 94.77 ± 0.48 | 96.35 ± 0.04 | +1.58 |
|  | GRU | 93.62 ± 0.91 | 96.16 ± 0.13 | +2.54 |
| DVS Gesture | SNN | 58.90 ± 3.68 | 64.20 ± 3.78 | +5.30 |
|  | SRNN | 49.72 ± 3.73 | 62.97 ± 7.79 | +13.25 |
|  | LSTM | 40.68 ± 15.57 | 62.42 ± 7.37 | +21.74 |
|  | GRU | 28.71 ± 11.81 | 66.21 ± 4.05 | +37.50 |

Time-series prediction provided a more selective test. Mean RMSE decreased in 9 of 12 task– backbone combinations, with reductions of approximately 10–11% in the strongest Mackey–Glass comparisons (Table 2). The remaining comparisons showed changes close to zero. Together, these results indicate that the computational benefit depends on the temporal prediction task and network architecture.

**Table 2:** Temporal-prediction RMSE and relative error reduction. Values are mean ± s.d. RMSE across *n* = 5 seeds per condition. Reduction is 100(RMSE_baseline_ −RMSE_BRIDGE_)*/*RMSE_baseline_; positive values indicate lower error and negative values indicate higher error.

| Task | Backbone | Baseline | BRIDGE | RMSE reduction |
|---|---|---|---|---|
|  |  | α = 0 | α = 0.5 | (%) |
| Lorenz | SNN | 8.7166 ± 0.7353 | 8.1972 ± 0.0161 | +5.96 |
|  | SRNN | 8.3626 ± 0.0148 | 8.1922 ± 0.0327 | +2.04 |
|  | LSTM | 7.8938 ± 0.0207 | 7.8531 ± 0.0206 | +0.52 |
|  | GRU | 8.0566 ± 0.0537 | 7.8722 ± 0.0231 | +2.29 |
| Mackey–Glass-17 | SNN | 0.2172 ± 0.0067 | 0.1937 ± 0.0064 | +10.82 |
|  | SRNN | 0.2149 ± 0.0178 | 0.1936 ± 0.0041 | +9.91 |
|  | LSTM | 0.1619 ± 0.0013 | 0.1621 ± 0.0042 | -0.12 |
|  | GRU | 0.1666 ± 0.0021 | 0.1657 ± 0.0021 | +0.54 |
| Mackey–Glass-30 | SNN | 0.2417 ± 0.0179 | 0.2379 ± 0.0155 | +1.57 |
|  | SRNN | 0.2413 ± 0.0146 | 0.2140 ± 0.0073 | +11.31 |
|  | LSTM | 0.1969 ± 0.0012 | 0.1973 ± 0.0009 | -0.20 |
|  | GRU | 0.1998 ± 0.0014 | 0.2004 ± 0.0010 | -0.30 |

The co-release fraction associated with the highest mean accuracy varied across tasks (Supplementary Fig. S8), complementing the fixed-*α* comparison with a task-specific parameter analysis.

## 3 Discussion

### 3.1 Resource asymmetry as a synaptic gain mechanism

BRIDGE links the resource dynamics of a cotransmitting terminal to the sign of its differential gain. A common presynaptic spike history drives the Glu and GABA branches, while their distinct depletion and recovery processes shape the net output. Sufficient asymmetry produces a nonmonotonic steady-state rate–drive curve: positive gain supports responsiveness at low activity, and negative gain produces a corrective change in drive above *f*_*g*_. Gain turnover can occur while the terminal remains net excitatory. This separation of differential gain from output polarity gives a quantitative account of how one presynaptic output unit can support both excitation and activity-dependent correction.

Short-term depression and differential E/I dynamics are established mechanisms of synaptic gain regulation [3, 20]. The present analysis identifies the turnover conditions for a commoninput, dual-resource architecture and connects its signed slope to recurrent input susceptibility. The resource variables also carry activity history: two inputs at the same current rate can produce different drive while their available resources differ. The steady-state curve specifies the eventual response, and resource recovery sets the path and time scale by which the synapse approaches it.

### 3.2 From differential gain to recurrent correction

The rate–resource analysis explains how the local synaptic rule enters recurrent feedback. At stable working points, negative differential gain lowers equilibrium input susceptibility relative to the local neuronal slope. The full Jacobian includes the recovery dynamics and determines whether perturbations decay. These complementary quantities distinguish the magnitude of the steady response from the time course of recovery.

The explicit recurrent networks demonstrate the corresponding local correction. Intrinsicdrive reductions elicit a shift toward excitation, and drive increases elicit a shift toward inhibition. Permanent output-sign reassignment tests this response under a persistent structural mismatch, with reduced activity burden relative to branchwise-matched common gain. Rate clamping further separates activity and balance outcomes: BRIDGE improves late E/I recovery relative to traditional and STD-only networks, while BRIDGE and common gain show similar aggregate E/I error. Across the tested perturbation protocols, the most consistent architecture-specific effects were directional local correction and activity recovery.

Larger circuits also converged on more similar late E/I ratios across realizations of the same perturbation. The standard deviation fell approximately as the inverse square root of network size in BRIDGE, and the two control architectures showed a similar decline. This pattern is consistent with population averaging as circuits grow. The recovery benefit of separate resources was evident at every size in the smaller firing-rate excursions relative to excitatory STD alone.

### 3.3 Biological implementation and computational use

The biological datasets constrain complementary parts of this mechanism. Hirai’s SuM→DG recordings show transmitter-specific short-term depression and frequency-dependent output filtering. The fitted model predicts the preceding gain transition. Meye’s branch-strength perturbations test the transfer of relative transmitter changes to postsynaptic output, while Pelkey’s interventions demonstrate biochemical branch addressability and shifts in output polarity. Pofahl’s measurements provide an inhibition-dominated DG working range for the recurrent sensitivity analysis. The finite-site simulations further quantify how release variability affects the reliability of signed synaptic output.

A direct test of the predicted gain transition is a dense frequency series measuring both transmitter branches and their combined postsynaptic effect. Accelerating Glu recovery should move *f*_*g*_ upward, and selective weakening of GABA should delay or abolish the turnover. Comparing responses to different input histories at the same final rate would additionally test the contribution of resource memory. A complementary computational comparison can hold the steady-state rate–drive curve fixed while varying the recovery dynamics, resolving how static gain and temporal feedback each contribute to network response.

The SNN experiments show that the same resource motif is compatible with feedforward, recurrent, and gated backbones, with task-dependent improvements in classification and temporal prediction. The *α* sweep identifies useful ranges for subsequent held-out optimization. Relating task performance to the activity regimes occupied by the trained units would connect these computational benefits more directly to their synaptic dynamics.

Asymmetric Glu/GABA resource dynamics thus provide a biologically motivated implementation of negative differential synaptic gain. By coupling opposing outputs to a shared activity history, a cotransmitting unit can generate bidirectional corrective drive and shape the response of recurrent networks to perturbations.

## 4 Methods

### 4.1 Model assumptions and notation

BRIDGE represents a cotransmitting presynaptic unit with two resource states driven by the same spike train {*t*_*k*_}. For branch *b* ∈ {e, i}, *x*_*b*_(*t*) ∈ [0, 1] is the available resource fraction, *U*_*b*_ ∈ [0, 1] is the effective depletion fraction per spike, and *τ*_*b*_ *>* 0 is the recovery time. The branches have separate depletion and recovery dynamics. Positive weights *A*_e_ and *A*_i_ combine release efficacy, quantal amplitude, and postsynaptic coupling. Both branches may depress; a non-depressing branch is obtained by setting its depletion fraction to zero.

We use discrete spike times for train responses and a continuous input rate *f* (*t*) for the rate model. Stationary frequency responses are derived separately for periodic and Poisson inputs. A common-gain control applies one resource state to both branch weights.

Subscripts e and i identify excitatory and inhibitory transmitter branches; uppercase E and I identify neuronal populations. *D* denotes effective drive and *I* denotes current. Subscripts n, s, and g denote net output, separate resources, and common gain, respectively. Frequencies are in hertz and time constants in seconds unless stated otherwise.

### 4.2 BRIDGE resource dynamics

Between spikes, each branch recovers toward full availability [4, 22, 25]:

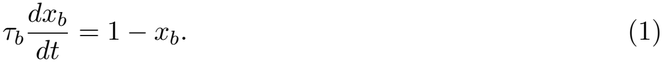

At spike time *t*_*k*_, the resource state changes from its pre-spike value 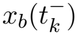 to its post-spike value 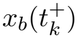:

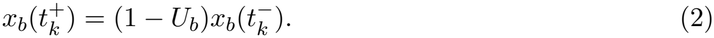

For periodic stimulation at frequency *f*, the steady-state pre-spike resource fraction *x*^*^(*f* ) is

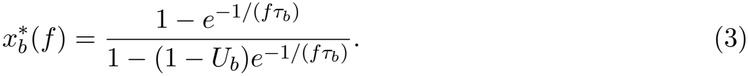

For *U*_*b*_ *>* 0, 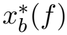 decreases from one toward zero as frequency increases. The static-resource limit is *U*_*b*_ = 0, 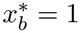. Here *U*_*b*_ describes effective population depletion.

### 4.3 Resource history and negative differential synaptic gain

For a time-varying rate *f* (*t*), resource availability obeys 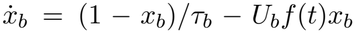. The instantaneous signed drive is

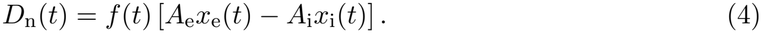

The states *x*_*b*_(*t*) retain the preceding input history. At constant input, their equilibrium defines the steady-state rate–drive relation.

For a stationary Poisson or continuous-rate input, the mean-field TM resource with utilization *U*_*b*_ and recovery time *τ*_*b*_ has the steady state

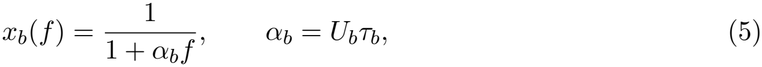

Here *α*_*b*_ = *U*_*b*_*τ*_*b*_ is the branch depression load, with units of time. For the gain-turnover analysis we assume *A*_e_ *> A*_i_ *>* 0, so the undepleted net drive is excitatory. The steady-state branch drives and their difference are

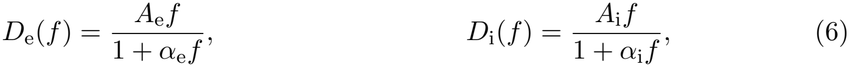

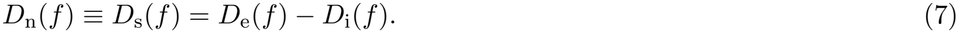

The differential gain is *g*(*f* ) = *dD*_n_(*f* )*/df* . The gain-turnover frequency *f*_*g*_ is a positive zero of *g*(*f* ) at which the slope changes from positive to negative. The net-drive zero-crossing frequency *f*_0_ is a positive solution of *D*_n_(*f*_0_) = 0.

#### Theorem 4.1

(Separate-resource gain turnover). *Under A*_e_ *> A*_i_ *>* 0 *and non-negative depression loads:*

1. *A positive finite gain-turnover frequency exists if and only if*

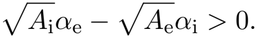

*When it exists*,

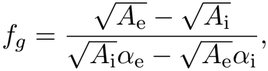
2. *An additional positive net-drive zero crossing exists if and only if*

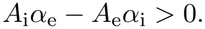

*When it exists*,

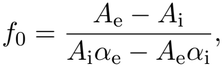
3. *The zero-crossing condition is strictly stronger than the gain-turnover condition. Whenever f*_0_ *exists*, 0 *< f*_*g*_ *< f*_0_.

*Proof*. For separate resources,

All denominators are positive. The sign of the derivative equals the sign of

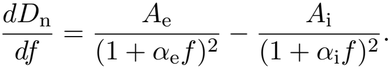

whose numerator is linear in *f* . Under the stated condition, that numerator starts positive, has negative slope, and crosses zero once, giving *f*_*g*_ and the claimed sign change. For the non-trivial zero crossing, cancelling the common factor *f >* 0 in *D*_n_(*f* ) = 0 likewise leaves a linear numerator; solving it gives *f*_0_ and the stated signs on either side. Writing *r* = (*A*_e_*/A*_i_)^1*/*2^ *>* 1, the zero-crossing condition is *α*_e_ *> r*^2^*α*_i_, which implies the gain-turnover condition *α*_e_ *> rα*_i_; direct subtraction gives

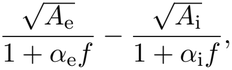

With other parameters fixed, increasing *U*_i_ shifts the onset of negative differential gain to higher rates as long as *α*_i_ *< α*_e_ *A*_i_*/A*_e_. When *f*_0_ exists, *f*_*g*_ *< f < f*_0_ is the regime of positive net drive with negative differential gain.

The theorem uses the stationary rate expression in Eq. 5. Periodic-train predictions use Eq.

3; agreement between their regime classifications is evaluated numerically.

### 4.4 Factorized common-gain control

The common-gain control uses one resource state with depression load *α* ≥ 0 for both branches:

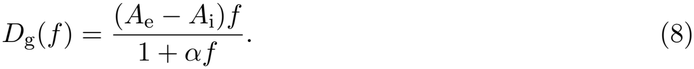

#### Proposition 4.1

(No turnover under a factorized common gain). *Under A*_e_ *> A*_i_ *>* 0 *and*

*α* ≥ 0,

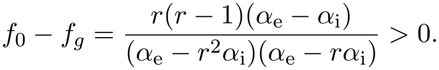

*Thus, the control can saturate but cannot produce a gain-turnover frequency*.

The control keeps the branch-amplitude ratio fixed throughout the input sweep.

### 4.5 Finite-site stochastic release and deterministic approximation

For finite-site release, branch *b* contains *N*_*b*_ sites with positive quantal amplitude *q*_*b*_. Let *p*_*b*_ ∈ [0, 1] be release probability conditional on availability; the effective probability at site *k* is *p*_*b*_*x*_*b,k*_. The net current is *I*_n_ = *I*_e_ − *I*_i_. The following results relate its mean and variability to deterministic branch output.

#### Proposition 4.2

(Mean correctness for any number of release sites). *Consider a presynaptic terminal with N*_e_ *glutamatergic and N*_i_ *GABAergic release sites. Let x*_*b,k*_ ∈ [0, 1] *be the specified pre-spike availability of site k in branch b, and let its release indicator have marginal distribution B*_*b,k*_ ∼ Bernoulli(*p*_*b*_*x*_*b,k*_). *Released quanta produce currents q*_e_ *and q*_i_. *Then, for* ***any*** *N*_e_, *N*_i_ ≥ 1:

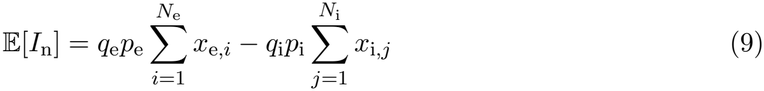

*By linearity of expectation, this equals the deterministic model prediction for every finite N*.

*Proof*. The total branch currents are 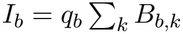. Since E[*B*_*b,k*_] = *p*_*b*_*x*_*b,k*_, linearity of expectation gives

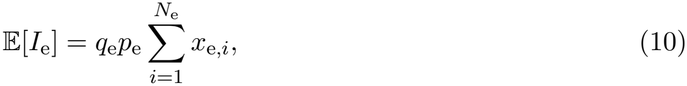

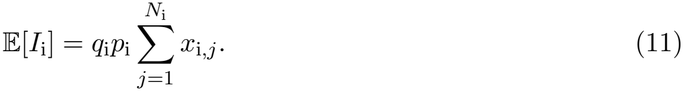

Therefore E[*I*_n_] = E[*I*_e_] −E[*I*_i_]. This holds for any *N*_e_, *N*_i_ ≥ 1, any specified branch availabilities, and arbitrary dependence among release indicators with the stated marginals. The variance formula below additionally assumes independent release indicators.

#### Proposition 4.3

(Cross-branch correlation changes variance, not mean). *Let K*_e_ *and K*_i_ *be the numbers of released Glu and GABA quanta on one event, with fixed marginal distributions and quantal amplitudes q*_e_, *q*_i_ *>* 0. *Then*

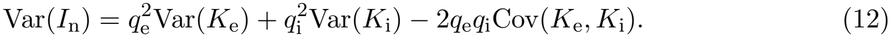

*Consequently, positive cross-branch covariance reduces net-current variance at fixed marginal release probabilities, while* E[*I*_n_] *is unchanged*.

*Proof*. Apply Var(*aX* − *bY* ) = *a*^2^Var(*X*) + *b*^2^Var(*Y* ) − 2*ab*Cov(*X, Y* ) to *I*_n_ = *q*_e_*K*_e_ − *q*_i_*K*_i_. Linearity of expectation establishes the second statement.

#### Proposition 4.4

(Variability scaling and distribution shape). *Under the same assumptions as Proposition 4*.*2, and assuming independence across release sites:*

1. *Suppose N*_e_ *and N*_i_ *grow proportionally and both branch variances grow linearly. If the mean net current per site stays bounded away from zero, then*

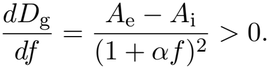

*Here C depends on release probabilities, quantal amplitudes, and E/I cancellation. Near exact cancellation, the CV is ill-conditioned and should not be used*.
2. *For finite N, I*_n_ *is a signed weighted sum of independent Bernoulli variables, a generalized Poisson-binomial distribution [26]*. *If all quantal amplitudes are equal, it is a shifted and scaled ordinary Poisson-binomial variable*.
3. *Let s*^2^(*N* ) = Var(*I*_n_) → ∞ *and assume that no single quantal weight dominates:*

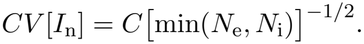

*Then the standardized net current converges to a Gaussian distribution by the Lindeberg– Feller central limit theorem*.

*Proof*. **(i)** Independence gives

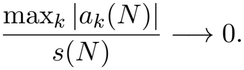

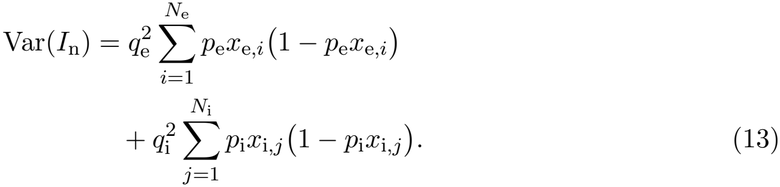

Under the stated proportional-growth and non-degeneracy assumptions this variance is Θ(*N* ), while the net mean is *Nµ* with |*µ*| *>* 0. Hence *CV* = Var(*I*_n_) */*|E[*I*_n_]| = *C/ N* . The prefactor *C* depends on branch statistics and increases as excitatory and inhibitory means cancel. At *µ* = 0, the net-current CV is undefined even though the standard deviation of the current normalized per site still decreases as *N* ^−1*/*2^.

i. Write the summands as *Y*_e,*k*_ = *q*_e_*B*_e,*k*_ and *Y*_i,*k*_ = −*q*_i_*B*_i,*k*_. Each is a two-point variable, so their convolution is a discrete signed weighted Bernoulli sum. Unequal positive and negative quantal weights require the generalized Poisson-binomial description; with equal weights, subtracting the inhibitory count gives a shifted and scaled ordinary Poisson-binomial variable.
ii. The centered summands are bounded by their quantal weights. Together, *s*^2^(*N* ) → ∞ and max_*k*_ |*a*_*k*_(*N* )|*/s*(*N* ) → 0 imply the Lindeberg condition, so (*I*_n_ − E*I*_n_)*/s*(*N* ) −→^*d*^ N(0, 1).

Finite-site simulations apply synaptic and membrane filters to stochastic and deterministic currents, then compare waveform error and output polarity (Fig. 3); site counts and parameter sampling are specified below.

### 4.6 Synapse-to-network rate model

Building on models of recurrent E/I networks [5, 6] and dynamic synapses [4], we couple population activity to the two BRIDGE resource variables:

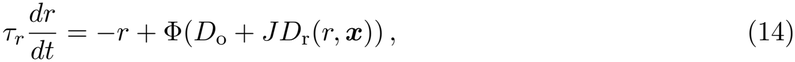

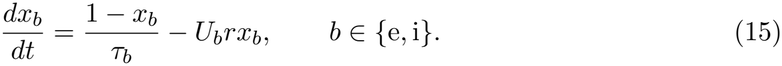

Here *r* is population firing rate, *τ*_*r*_ is its relaxation time, ***x*** = (*x*_e_, *x*_i_), *D*_o_ is external drive, and *J* ≥ 0 is recurrent coupling. BRIDGE supplies *D*_r_(*r*, ***x***) = *r*(*A*_e_*x*_e_ − *A*_i_*x*_i_). The neuronal response function is Φ(*z*) = *r*_max_*/*[1 + exp(−*βz*)], with maximum rate *r*_max_ and slope parameter *β*.

Static, common-gain, and separate-resource models are compared under matched-rate and fixed-input conditions. At each equilibrium, local stability requires max Re(*λ*) *<* 0 for the full rate–resource Jacobian. Numerical parameters and operating-point sweeps are specified below.

#### Proposition 4.5

(Equilibrium input susceptibility). *Consider Eqs. 14–15, with J*≥ 0 *and* Φ *continuously differentiable near a fixed point r*^*^. *Let*

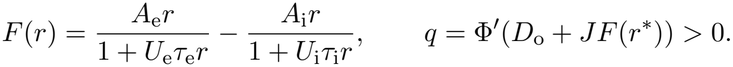

*If* 1 − *JqF* ^′^(*r*^*^) ≠ 0, *the local equilibrium branch has input susceptibility*

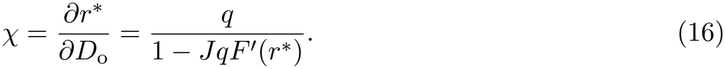

*For F* ^′^(*r*^*^) *<* 0, *the normalized susceptibility satisfies* 0 *< χ/q* ≤ 1, *with strict inequality when*

*J >* 0.

*Proof*. At equilibrium, 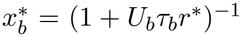 and *r*^*^ = Φ(*D*_o_ + *JF* (*r*^*^)). Implicit differentiation gives *χ* = *q*[1 + *JF* ^′^(*r*^*^)*χ*], yielding Eq. 16 and the stated inequalities.

On a stable branch, *χ* measures the equilibrium rate response to external input and *q* is the local neuronal slope. Transient stability uses the full Jacobian given in Supplementary Theory (p. 38).

## 5 Experimental procedures and outcome measures

### 5.1 Measures of synaptic output, activity recovery, and E/I balance

Synaptic gain is *g*(*f* ) = *dD*_n_*/df* . Numerical turnover detection identifies its first positive-to-negative crossing, *f*_*g*_; the first positive zero of net drive is recorded separately as *f*_0_.

Finite-site simulations compared the filtered stochastic waveform *y*_*s*_ with its deterministic counterpart *y*_*d*_. With *µ*_e_ and *µ*_i_ denoting the deterministic branch amplitudes, waveform error was

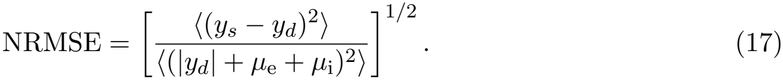

Polarity disagreement was the fraction of release events for which stochastic and deterministic net amplitudes had opposite signs. Relative mean bias was ⟨*y*_*s*_ −*y*_*d*_⟩*/* max(|⟨*y*_*d*_⟩|, *µ*_e_ +*µ*_i_). Before analysis, functional equivalence was defined as NRMSE *<* 0.10 and polarity disagreement *<* 0.05 for the same parameter–frequency combination.

For recurrent networks, *E*(*t*) and *I*(*t*) denote the magnitudes of recurrent glutamatergic and GABAergic input in each 20-ms bin. Their bounded balance index was

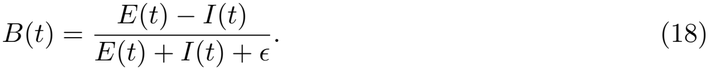

Each network’s pre-intervention rate *r*_0_ and balance *B*_0_ define its working point. Over the post-intervention interval [*t*_1_, *t*_2_], excess and suppressed activity are

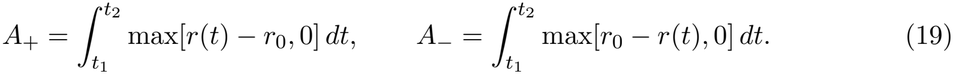

The primary outcome is *A*_−_ after E-to-I reassignment and *A*_+_ after I-to-E reassignment. With subscript l denoting the late evaluation window, late rate and E/I errors are |*r*_l_ − *r*_0_|*/r*_0_ and

|*B*_l_ − *B*_0_|. Joint recovery requires errors no greater than 0.20 and 0.05, respectively. For the co-release component, *D*_c_ = *E*_c_ − *I*_c_. Directional correction after an intrinsic-drive step Δ*u* is *C*_d_ = − sgn(Δ*u*)Δ*D*_c_, where Δ*D*_c_ is the change from baseline. Positive *C*_d_ indicates a shift opposing the imposed activity change.

Classification effects are mean accuracy differences in percentage points. Prediction effects are 100(RMSE_0_ −RMSE_*α*_)*/*RMSE_0_, where subscripts denote the baseline and co-release models; positive values indicate improvement. Unless stated otherwise, tests are two-sided, intervals are percentile-bootstrap 95% confidence intervals, and Holm correction is applied within each pre-specified family. Seed-matched simulations are paired; biological resampling units are specified below.

### 5.2 Numerical validation of gain turnover and finite-site release

Thresholds from Theorem 4.1 were checked against numerical roots for 800 fixed-seed parameter draws with positive branch amplitudes and non-negative depression loads. A separate 672-condition grid compared the rate and periodic-train classifications, with agreement of at least 90% set as the acceptance criterion. Three hundred common-gain draws tested Proposition 4.1. Periodic, Poisson, and bursty gamma-renewal inputs were evaluated over 0.2–60 Hz; random conditions used 20 fixed seeds and 8,000 events per rate. Finite-site threshold recovery used 150 repetitions at *N* = 10, 20, 50, 100 and 800 events per frequency. Turnover detection required a second-order Savitzky–Golay-smoothed derivative crossing and at least a 10% post-peak decline; mean-drive estimates remained unsmoothed. For cross-branch correlation, both marginal binomial distributions were held fixed while feasible correlation was varied; net-current standard deviation was normalized to independent release.

### 5.3 Electrophysiological data and parameter estimation

Hirai et al. [15] provided glutamate/GABA cotransmission recordings from supramammillary projections to dentate granule cells (SuM→DG). Their Figure 1G reports paired-pulse ratios of 0.46 ± 0.03 for Glu and 0.60 ± 0.03 for GABA at a 100-ms interval (mean ± SEM, *n* = 8 per component).

EPSC trains at 2.5 mM Ca^2+^ were fitted over stimuli 2–10 at 5, 10, and 20 Hz (*n* = 27 observations), using reported SEMs in a weighted least-squares Gaussian likelihood. Stimulus 1 defines the 100% normalization and was excluded. The fitted response was *R*_*n*_ = *b* +(100 − *b*)*x*_*n*_, where *x*_*n*_ is the pre-spike resource fraction obtained from Eqs. 1–2, with *x*_1_ = 1. Parameters were depletion fraction *U*, recovery time *τ*_x_, and residual percentage *b*. We propagated two interpretations of *b*: a measurement floor excluded from current dynamics, or a non-depressing component with normalized response *b/*100 + (1 − *b/*100)*x*(*f* ). A zero-residual fit served as a sensitivity control. Three hundred fixed-seed parametric bootstrap resamples refitted these models. Fixed-amplitude thresholds used *A*_e_ = 1.005 and *A*_i_ = 0.5 for illustrative sensitivity analysis.

Functional branch weights were estimated from the cell-level spike data in Hirai et al. Figure 6 using

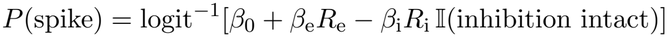

Here *R*_e_ and *R*_i_ are branch responses, *β*_0_ is the intercept, *β*_e_ and *β*_i_ are fitted weights, and I is one when inhibition is intact and zero under picrotoxin. Light and picrotoxin observations were fitted jointly and resampled as cell pairs within frequency. Glu and GABA train cells were resampled independently within branch and frequency because their cross-branch pairing is unavailable. Five hundred bootstrap fits yielded *ρ* = *β*_i_*/β*_e_. Independent samples from this weight distribution and the 300 resource fits generated 5,000 joint draws per residual interpretation. A turnover required the exact periodic-model slope to cross from positive to negative over 0.25–100 Hz.

For each of the three residual-response interpretations, 5,000 joint draws were also evaluated at 5 Hz for marginal gain and the signs of drive changes under symmetric rate perturbations. Recurrent-network sensitivity used balance targets *B*_0_ = −0.40, −0.52, −0.60, −0.70, −0.75, with four architectures, both reassignment directions, and 12 new seed-matched networks per target. The final two targets overlap the inhibition-dominated range informed by Pofahl et al. [24], who reported inhibition 5.66–7.50 times excitation in medial-perforant-path recordings.

### 5.4 Cross-condition and cross-pathway biological prediction

Figure-level source workbooks from Hirai et al. [15], Meye et al. [13], and Pelkey et al. [14] were converted to a long-format table retaining workbook, worksheet, row, column, and SHA-256 hash. Original workbooks were preserved. Analyses used recorded cells as observational units because cell-to-animal mappings were unavailable.

For Hirai et al., cell-level EPSC and IPSC trains at 2.5 mM Ca^2+^ were summarized over late pulses 8–10. Differential frequency scaling was tested directly with late response ∼ branch × log_2_(*f/*5) using one late-train average per cell and HC3 heteroscedasticity-robust standard errors. The branch-by-frequency interaction quantifies the difference between Glu and GABA slopes; the GABA slope itself was estimated with its confidence interval. Their Figure 6 spike probabilities were paired by source-cell identifier across control sinusoidal current, paired SuM light stimulation, and light stimulation with picrotoxin. Early and late epochs were defined before testing as pulses 1–3 and 8–10. Paired condition contrasts were Holm-corrected across the 12 frequency–epoch–contrast combinations. To assess predictive transport across frequencies, logistic output mappings were fitted on two frequencies and evaluated on the third. We compared an excitation-only model, a model with excitation plus an intact-inhibition condition indicator, and a mechanistic E+I model in which the measured GABA trace contributes only when inhibition is intact. Synaptic train parameters were estimated from their Figure 5 independently of the Figure 6 spike data.

For Meye et al., the prespecified cell-level contrasts were saline-control versus cocaine-control (perturbation) and cocaine-VGAT versus cocaine-control (rescue) for the GABA_*A*_R/AMPAR ratio, spike fidelity relative to baseline, and IPSC recovery after the third train. We report mean differences, percentile-bootstrap 95% intervals, Hedges’ *g*, and two-sided Welch tests with Holm correction across the six contrasts; exact permutation tests with the same correction were used as a small-sample robustness check. A separate cross-condition prediction withheld the complete cocaine+ACSF 5-Hz, ten-pulse spike-fidelity train until final scoring. Saline picrotoxin and ACSF means defined the pulse-specific inhibitory template

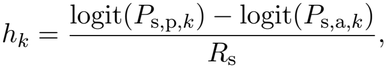

Here *k* indexes the pulse, and *R*_s_ and *R*_c_ are the independently measured saline and cocaine GABA_*A*_R/AMPAR ratios. For spike probability *P*, the first condition label denotes saline (s) or cocaine (c), and the second denotes picrotoxin (p) or ACSF (a). The predicted cocaine response was logit^−1^{logit(*P*_c,p,*k*_) − *R*_c_*h*_*k*_}. Excitation-only and unchanged-ratio predictions were fixed before scoring. Cells were resampled independently within each pulse condition and ratio group for 5,000 bootstrap draws; the held-out cells entered only the scoring distribution.

For Pelkey et al., direct inhibition of glutamic acid decarboxylase with semicarbazide was used as the discovery perturbation, and dietary vitamin-B6 deficiency was retained as a mechanistically related cross-perturbation holdout. For each perturbation, the predicted directions were lower GABA conductance, higher or unmasked glutamate conductance, and a lower GABA/Glu ratio. We report cell-level medians, bootstrap 95% intervals for median differences, Cliff’s *δ*, and two-sided Mann–Whitney tests with Holm correction across the six planned comparisons. In the additional polarity prediction, each cell’s measured conductances were mapped without fitted coefficients to *D*(*V* ) = *g*_e_(*E*_e_ − *V* ) − *g*_i_(*V* − *E*_i_) at the fixed values *V* = −52 mV, *E*_e_ = 0 mV, and *E*_i_ = −70 mV. This design provides a held-out cross-perturbation test of output direction. The source article reports 4 control and 4 semicarbazide mice, and 4 B6-sufficient and 9 B6-deficient mice; because the workbook does not map cells to individual mice, inference is conducted at the cell level. The inverse-hyperbolic-sine scale used for the two ratio panels is a visualization transform that accommodates zero values and is excluded from the tests.

### 5.5 Release correlations and input-count approximations

Within-channel release correlation was varied over *ρ* = 0–0.4 for 20–40 sites. Binomial input counts were compared with Gaussian approximations at *N* = 10^3^, 5×10^3^, and 10^4^ inputs (Supplementary Fig. S1).

### 5.6 Stochastic release across finite site counts

Finite-site simulations used *N* ∈ {10, 20, 30, 40, 50} independent sites per branch and 100 parameter sets sampled at fixed intervals from the 300 resource bootstrap fits. At 5, 10, and 20 Hz, Glu and GABA counts were sampled independently from binomial distributions. We used the fitted Glu dynamics and treated the GABA resource as static (*U*_i_ = 0, *x*_i,*j*_ = 1) because late GABA responses varied little across 5–20 Hz (Fig. 2b). This approximation concerns frequency dependence late in the train. At 5 and 20 Hz, the measured late GABA responses averaged 25.4% and 23.4% of the first response, respectively. Both stochastic and deterministic currents passed through exponential synaptic (*τ*_s_ = 5 ms) and membrane (*τ*_*m*_ = 20 ms) filters. Waveform and polarity measures were evaluated after a 100-spike burn-in. Figure 3a,b used the boot-strap draw nearest the joint median (*U* = 0.8572, *τ*_x_ = 1107.5 ms), 10-Hz stimulation, and 80 fixed-seed realizations per displayed *N* .

### 5.7 Matched recurrent networks under permanent E/I perturbations

#### Matched networks and permanent identity reassignment

A recurrent network of *N* = 500 LIF neurons (E:I = 3:1, fixed out-degree *K* = 50; 50% of baseline E neurons assigned to the corelease branch where applicable, equivalent to 37.5% of the complete network) was simulated for 6 s with *dt* = 0.5 ms. Four main conditions used identical connectivity, conversion masks, and noise seeds: static traditional E/I, excitatory STD without co-release, spike-locked common-gain cotransmission, and spike-locked branch-resolved BRIDGE cotransmission. In the common-gain control, one resource variable multiplied both co-released branches. Its Glu and GABA event amplitudes were matched exactly to BRIDGE at the 5-Hz periodic steady state, so the comparison changed only whether the two branches could move relative to one another. Before any conversion, inhibitory gain and background drive were calibrated using baseline-only networks and then refined using only the 1–2-s baseline windows of the validation cohort. The required cohort-mean tolerances were 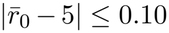 Hz and 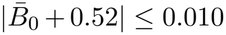, using the balance index in Eq. 18. The reported estimand and calibration estimand were identical: *B*(*t*) was evaluated in each 20-ms bin before averaging. Parameters remained fixed throughout the post-conversion interval.

We used an in-silico stress test of persistent presynaptic output-sign mismatch. At 2 s, a fixed fraction of neurons underwent permanent presynaptic output-identity reassignment. In the E-to-I protocol, randomly selected ordinary E neurons thereafter emitted conventional inhibitory output; in the I-to-E protocol, selected I neurons thereafter emitted conventional excitatory output; and in the balanced-swap protocol, equal numbers were reassigned in both directions while preserving the global cell counts. Incoming connectivity, intrinsic dynamics, targets, and presy-naptic weight heterogeneity were unchanged. Primary conversion masks targeted conventional E and I neurons, preserving the co-release population during the perturbation. Newly excitatory outputs in STD-containing conditions were initialized at the exact 5-Hz periodic steady-state resource value to avoid an artificial fully recovered transient.

Eight pilot networks, disjoint from calibration and validation, evaluated conversion fractions of 1, 2, 4, 6, and 8% of all neurons using only the traditional architecture. A fraction was eligible when E-to-I produced a measurable nonsilent reduction and I-to-E produced a measurable nonsaturated increase; no architecture contrast entered selection. The default middle level, 4%, passed these checks and was frozen as primary. Thirty new seed-matched networks supplied the primary validation outcomes. Outcomes over the permanent 2–6-s conversion interval used the protocol-specific activity burdens in Eq. 19; the late state was averaged over 5–6 s. BRIDGE was compared with the branchwise-matched common-gain control for activity burden and late E/I error in each direction using paired two-sided Wilcoxon tests with direction-family Holm correction.

#### Rate-only decoupling control

To separate changes in firing rate from changes in recurrent E/I balance, a proportional–integral rate controller was applied equally to all four architectures after conversion. The controller used population firing rate as its sole feedback signal and targeted each trial’s own 1–2-s baseline rate. Controller gains were frozen using eight traditional-E/I pilot networks and a rate-only score before the 30 validation seeds were run. The prespecified quality criterion required median late-rate error below 10% in every architecture and conversion direction. Late E/I error was then compared between BRIDGE and each control with Holm correction across the six architecture–direction tests.

#### Intrinsic-drive perturbation

The smallest symmetric intrinsic-drive steps producing non-silent, nonsaturated responses in traditional-E/I pilot networks were selected and fixed at |Δ*u*| = 0.5. Thirty new seed-matched common-gain and BRIDGE networks received each step with output identities unchanged. The primary outcome was *C*_d_; the two direction-specific paired tests were Holm-corrected.

#### Co-release-prevalence sensitivity

The 50%-of-E reference prevalence was then varied from 0% to 90% of E neurons in 10-percentage-point increments at the frozen 4% structural conversion. The zero-prevalence endpoint reduces to the STD-only architecture. Each prevalence received an independent baseline-only calibration on 16 networks and validation-cohort refinement on 30 new networks, all disjoint from the main validation seeds; the same 5-Hz and *B*_0_ = −0.52 tolerances were enforced before E-to-I or I-to-E outcomes were evaluated. The sweep stops at 90% because co-release neurons remain excluded from the conversion mask and a 4%-of-network ordinary-E cohort must remain available. Prevalence effects are summarized by median and interquartile trajectories across the tested fractions.

#### Explicit network-size scaling

The frozen 4% E-to-I and I-to-E protocols were repeated in explicit sparse networks at *N* = 100, 200, 500, 1000, 2000, 5000, and 10,000 (*n* = 12 seed-matched networks per condition and size). Out-degree remained fixed at *K* = 50, and every size–architecture combination was matched using only its own pre-conversion baseline window before outcomes were evaluated. All neurons and recurrent connections were simulated explicitly. Scale persistence was summarized as the seed-paired percentage reduction in the appropriate activity burden relative to STD-only (Fig. 5f). The corresponding finite-size E/I-ratio dispersion is reported in Fig. 5g over the same *N* = 100–10^4^ range.

The compact parameter labels denote resting voltage *V*_0_, reset voltage *V*_r_, threshold voltage *V*_t_, synaptic time constant *τ*_s_, refractory period *τ*_f_, and resource-recovery time *τ*_x_. The shared network parameters are reported in Table 3, following reproducible network-description recommendations [27]. Unless explicitly varied, the same values were used across conditions.

**Table 3:**
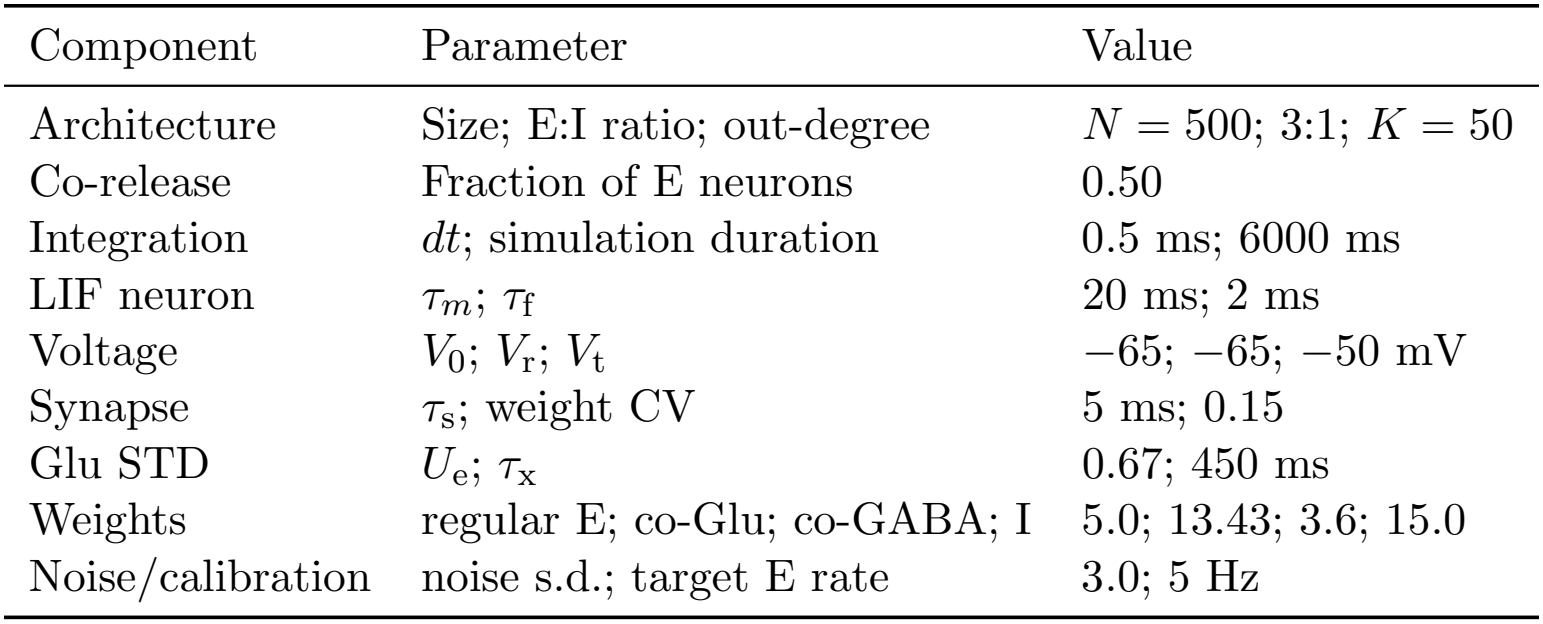
Shared parameters for the recurrent structural-compensation experiments.

| Component | Parameter | Value |
|---|---|---|
| Architecture | Size; E:I ratio; out-degree | *N* = 500; 3:1; *K* = 50 |
| Co-release | Fraction of E neurons | 0.50 |
| Integration | *dt*; simulation duration | 0.5 ms; 6000 ms |
| LIF neuron | τ<sub>*m*</sub>; τ<sub>f</sub> | 20 ms; 2 ms |
| Voltage | *V*<sub>0</sub>; *V*<sub>r</sub>; *V*<sub>t</sub> | −65; −65; −50 mV |
| Synapse | τ<sub>s</sub>; weight CV | 5 ms; 0.15 |
| Glu STD | *U*<sub>e</sub>; τ<sub>x</sub> | 0.67; 450 ms |
| Weights | regular E; co-Glu; co-GABA; I | 5.0; 13.43; 3.6; 15.0 |
| Noise/calibration | noise s.d.; target E rate | 3.0; 5 Hz |

### 5.8 Resource-architecture and postsynaptic-conductance controls

#### Biological phase map

The conditional gain-turnover frequency *f*_*g*_ was calculated on a grid of *U*_e_ = 0.15–0.95 and *τ*_e_ = 200–1800 ms while holding the illustrative mapping fixed at *w*_e_ = 1.5 and *w*_i_ = 0.5. The numerically estimated boundary for a crossing over 0.25–100 Hz and two reference parameterizations were overlaid. The reference markers are (*U*_e_, *τ*_e_) = (0.6587, 511.1 ms) and (0.8574, 1085.2 ms); bootstrap uncertainty is reported separately in Supplementary Fig. S2. The map reports sensitivity of the turnover frequency to resource parameters at the stated fixed branch weights.

#### Matched reduced-model controls

The rate model used *τ*_*r*_ = 20 ms, *r*_max_ = 100 Hz, and *β* = 0.10. The iso-rate family adjusted external input at each *J* to maintain 5 Hz in the static, common-gain, and separate-resource models. The fixed-input family calibrated a common input at *J* = 0, then held it constant while enumerating equilibria over *J* = 0–20. A working-rate sweep fixed *J* = 8 and adjusted input to match rates from 0.25 to 30 Hz (Fig. 4a,b). Equilibria were classified by Jacobian eigenvalues; fixed-point residuals below 10^−7^ and recovery of the 5-Hz root at *J* = 0 were required checks.

#### Conductance-based robustness

A single-compartment LIF neuron used *E*_*l*_ = *V*_r_ = −65 mV, *V*_t_ = −50 mV, *τ*_*m*_ = 20 ms, a 2-ms refractory period, and excitatory reversal 0 mV. An 81 × 81 grid varied inhibitory reversal *E*_i_ from −85 to −45 mV and the GABA/Glu conductance scale from 0.20 to 2.20. External current was calibrated to 5 Hz. Firing was evaluated across presynaptic rates to locate marginal-gain zero crossings. BRIDGE used frequency-dependent Glu depletion and static GABA resources; the common-gain control scaled both conductances by one state. The primary domain comprised 5,751 inhibitory/shunting points with *E*_i_ ≤ *V*_t_; above-threshold points were marked separately.

### 5.9 Classification and temporal prediction with co-release SNNs

Four backbones were evaluated with co-release fractions *α* = 0 and 0.5: a feedforward SNN, a recurrent SNN (SRNN), and recurrent SNNs with LSTM or GRU gating. Here *α* is the fraction of excitatory units with a co-release branch, distinct from the branch depression load *α*_*b*_. The SNN implementation retained a dynamic resource only on the Glu branch. Each spike from a cotransmitting unit recruited the available Glu resource and drove a GABA output proportional to its synaptic weight. These branch currents were combined before LIF integration. Training used surrogate-gradient optimization [28].

Classification covered P-SMNIST and S-MNIST derived from MNIST [29], the SHD and SSC auditory event datasets [30], MNIST and Fashion-MNIST [29, 31], N-MNIST [32], and DVS Gesture [33]. Continuous prediction covered the Lorenz system [34] and Mackey–Glass series with delays 17 and 30 [35]. The supplied classification summaries contain four seeds for SNN and SRNN and five seeds for the LSTM- and GRU-gated variants; all prediction summaries contain five seeds. The SRNN *α* sweep used *α* ∈ {0, 0.1, 0.25, 0.5, 0.75, 1}, with four seeds at *α* = 0 and 0.5 and five at the other settings.

Tables 1 and 2 report the supplied architecture–task means and standard deviations. Classification gain is co-release minus baseline accuracy in percentage points; prediction improvement is the relative mean RMSE reduction, 100(RMSE_baseline_ −RMSE_BRIDGE_)*/*RMSE_baseline_. Supplementary Fig. S8 retains the task-specific *α* sweep. Its bars show supplied means, whiskers show one supplied standard deviation, and diamonds show the supplied best run. The best tested *α* denotes the grid value with the highest supplied mean accuracy.

## Acknowledgments

This work was supported by the Major Program of Xiangjiang Laboratory (No. 24XJJCYJ01001), the National Science and Technology Major Project “Brain Science and Brain-Inspired Intel-ligence” (2025ZD0215700), and the National Natural Science Foundation of China (72025405, 72421002, 92467302, 72088101, 72301285, 72474223).

## Supplementary Results

### Release correlations delimit the independent-site approximation

**Figure S1:**
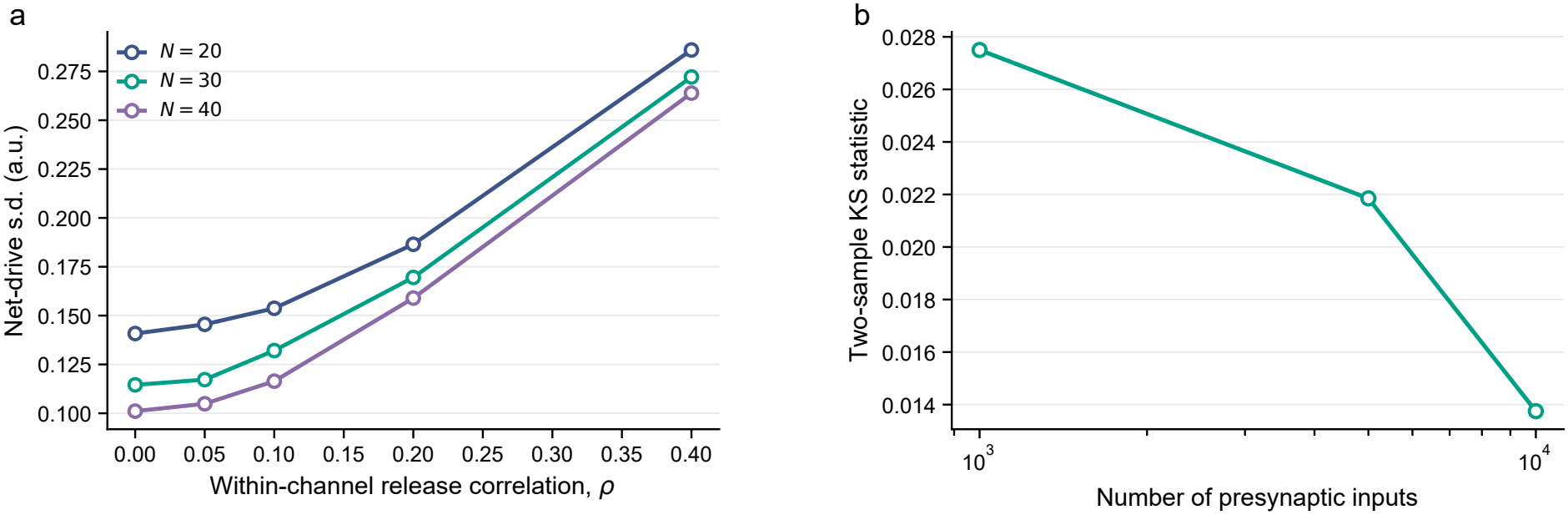
Release correlations and input-count approximations. (a) Increasing within-channel release correlation raises the standard deviation of net drive for 20–40 release sites, delimiting the independent-site regime used by the mean-field abstraction. (b) Exact binomial input counts and the Gaussian approximation become increasingly similar as the number of presynaptic inputs grows. The agreement supports the Gaussian approximation for aggregated presynaptic input counts.

**Figure S2:**
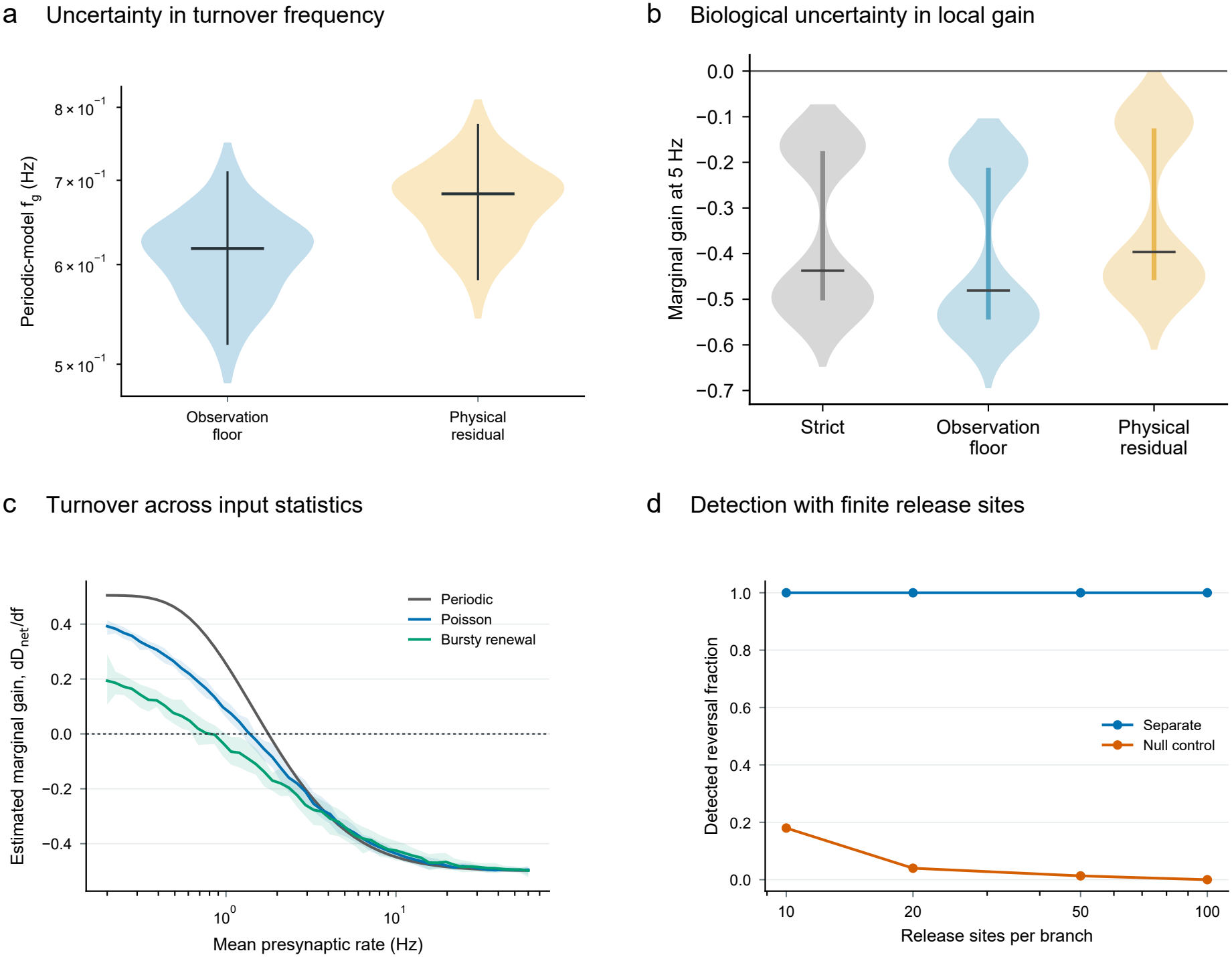
Gain turnover survives biological uncertainty and variable input statistics. (a) Periodic-model *f*_*g*_ over 300 bootstrap refits for the observation-floor and physical-residual interpretations at fixed effective branch amplitudes; bars show medians and 95% ranges. (b) Marginal gain at 5 Hz across 5,000 joint Hirai bootstrap draws per interpretation, including uncertainty in effective branch weights. Negative gain occurs in all 5,000 strict-model and observation-floor draws and in 4,999 of 5,000 physical-residual draws; bars show medians and interquartile ranges. Panels a and b propagate different uncertainty sets and report different outputs. (c) Marginal-gain curves under periodic, Poisson, and bursty renewal input; lines show seed means and shading shows 95% ranges for random inputs. (d) Fraction of 150 finite-site repetitions classified as showing a material gain turnover. Separate resources are detected in all repetitions; false detections for the exact-monotone common-gain null decline to zero by *N* = 100.

### Gain turnover persists across input statistics and model uncertainty

Allowing a residual response improved the train fit over the strict zero-residual control (ΔAICc = 115.4). The observation-floor and physical-residual interpretations share one likelihood but imply different current equations. Both retain gain turnover under parameter uncertainty (Supplementary Fig. S2a,b) and variable input timing (panel c). Finite-site detection remains reliable for separate resources, whereas small-*N* false detections in the common-gain null decline with site number (panel d).

### Resource asymmetry limits recurrent amplification across model conditions

The parameter map identifies the presynaptic release and recovery combinations that support gain turnover. Reduced recurrent models then test the consequence of this turnover under matched-rate and fixed-input conditions. Finally, a conductance-based model delineates the postsynaptic conditions under which the same output signature persists.

**Figure S3:**
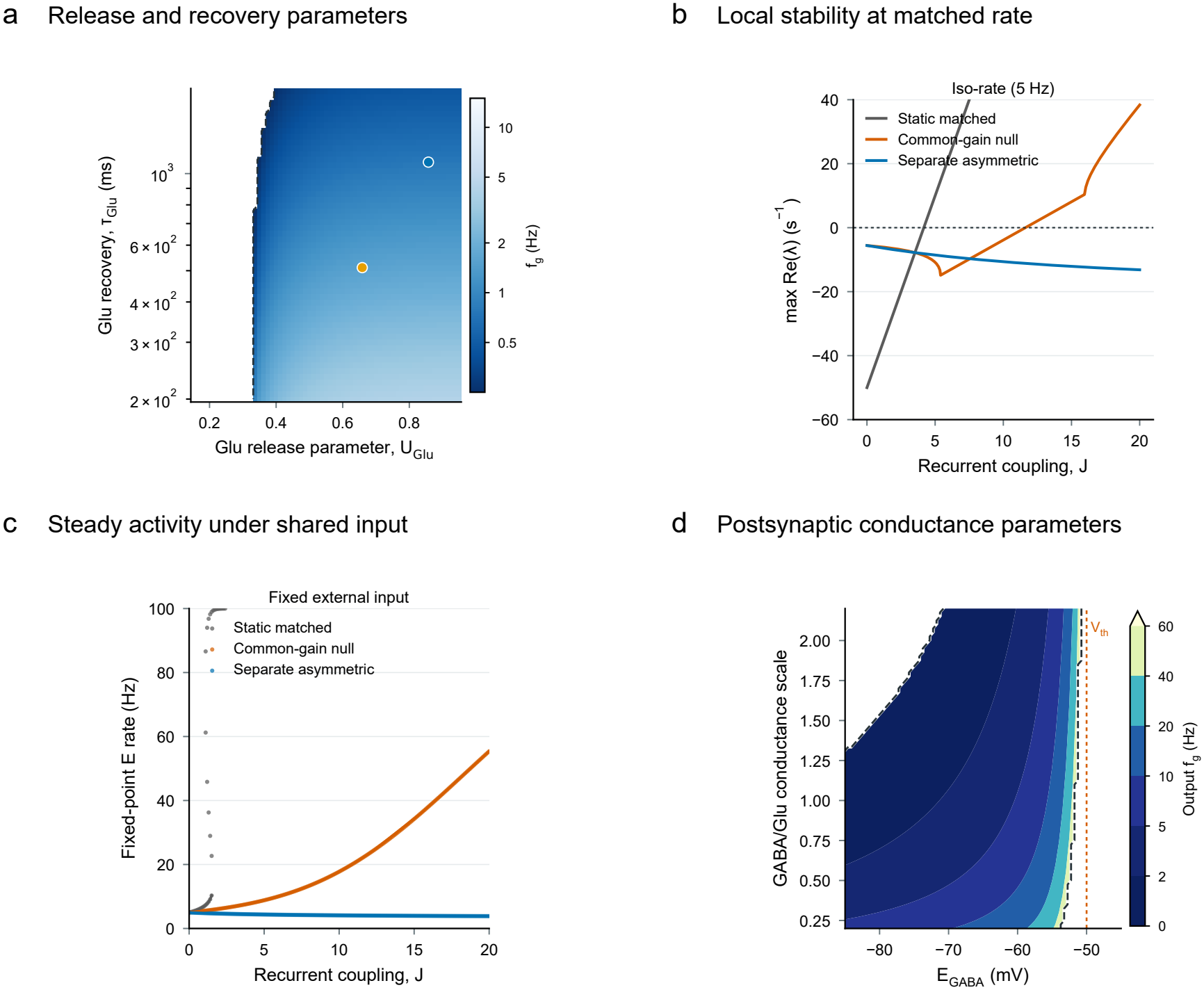
Presynaptic and postsynaptic parameters delimit resource-dependent gain control. (a) Conditional *f*_*g*_ map over Glu release and recovery parameters, with two reference parameterizations overlaid (gold and blue markers, respectively, as defined in Methods). The map varies resource parameters at fixed effective branch amplitudes. (b) Dominant Jacobian eigenvalue with external input recalibrated at every recurrent coupling *J* to maintain a 5-Hz operating rate. Values below zero indicate local stability. (c) Fixed-point firing rates under one common external input calibrated at *J* = 0, showing how steady activity changes as recurrent coupling increases. (d) Conductance-based output *f*_*g*_ over GABA reversal potential and the GABA/Glu conductance scale. White regions indicate no crossing within the tested range; the region above spike threshold is shown as a sensitivity region.

### Independent biological perturbations shift transmitter balance and output

Meye’s data show that VGAT expression reverses changes in inhibitory branch strength and output associated with cocaine (Supplementary Fig. S4a,b). Pelkey’s enzyme inhibition and B6-deficiency perturbations reduce the GABA/Glu ratio (panels c,d), supporting selective biochemical control of transmitter-branch strength. For Hirai, prediction at held-out frequencies reduced pooled RMSE from 18.6 to 16.5 percentage points when measured E and I traces replaced excitation alone. An intact-inhibition indicator reached 16.8 points, showing that detailed GABA dynamics add limited predictive information beyond the presence of inhibition.

**Figure S4:**
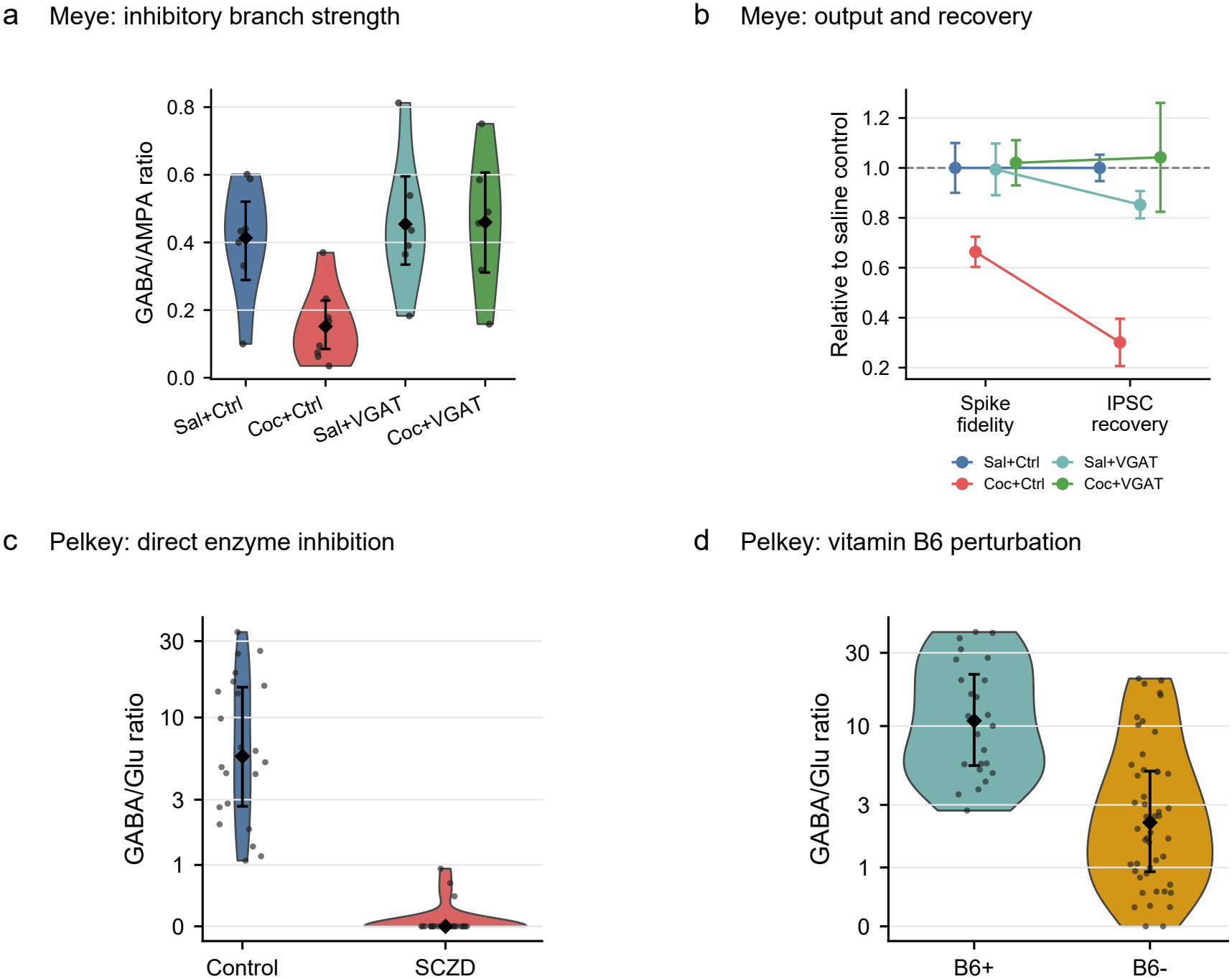
Branch-selective interventions alter synaptic balance and output. (a) Meye et al. [13]: GABA_*A*_R/AMPAR ratios after saline or cocaine treatment with control or VGAT virus (*n* = 6–8 cells/group); black symbols show means and 95% bootstrap intervals. (b) Spike fidelity and IPSC recovery relative to saline controls (means ± SEM; *n* = 5–6 cells/group). (c,d) Pelkey et al. [14]: GABA/Glu ratios after semicarbazide treatment (22 control, 23 treated cells) or B6 deficiency (24 sufficient, 48 deficient cells). Black diamonds show medians and interquartile ranges; inverse-hyperbolic-sine axes retain raw-value tick labels. Violin envelopes show distributions and dots show cells. Pelkey reports 4/4 and 4/9 mice, respectively. The source data provide cell-level observations with unassigned animal membership; cells are the analysis units.

### Activity compensation persists across inhibition-dominated working points

Pofahl et al. measured medial-perforant-path-evoked excitation and inhibition in dentate granule cells in vivo [24]. Their inhibition-to-excitation conductance ratio corresponds to a balance interval of approximately *B*_0_ = −0.765 to −0.700, shown by the shaded region in Supplementary Fig. S5. These medial-perforant-path data define an externally motivated, strongly inhibition-dominated DG working region for the recurrent sensitivity analysis. The four panels compare activity burden and late E/I error across the same baseline-matched networks, separating activity compensation from recovery of aggregate balance.

**Figure S5:**
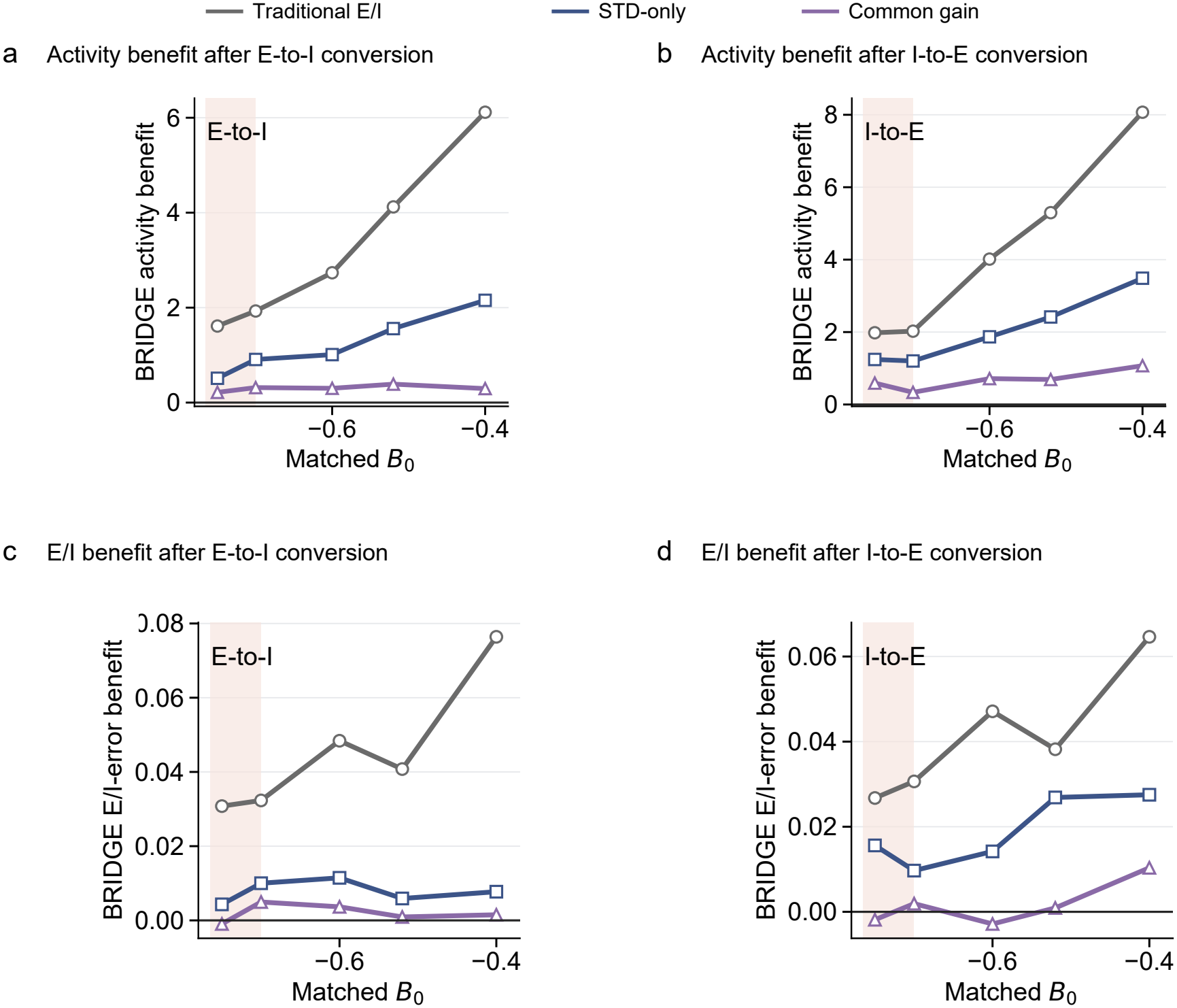
Activity and E/I outcomes across matched recurrent working points. (a,b) Median BRIDGE activity-burden benefit relative to traditional E/I, STD-only, and branchwise-matched common gain after E-to-I and I-to-E conversion, respectively. (c,d) Corresponding median benefit in late recurrent E/I error. Positive values favour BRIDGE; shaded bands mark the DG interval derived from the conductance ratio reported by Pofahl et al. [24]. At the most inhibition-dominated target, the late E/I-error comparison slightly favors common gain. Each point contains 12 seed-matched networks after target-specific baseline calibration.

### Biological uncertainty in the predicted gain-turnover frequency

A paired-cell output model estimated a positive inhibitory contribution, with a bootstrap median *β*_i_*/β*_e_ ratio of 0.501 (95% interval 0.164–0.645). Joint sampling of the functional weights and dynamical fits yielded gain-turnover frequency distributions for the strict-TM, observation-floor, and physical-residual interpretations (Supplementary Fig. S6). The measured 5–20-Hz trains constrain the high-frequency responses, while the periodic model estimates the preceding turnover.

**Figure S6:**
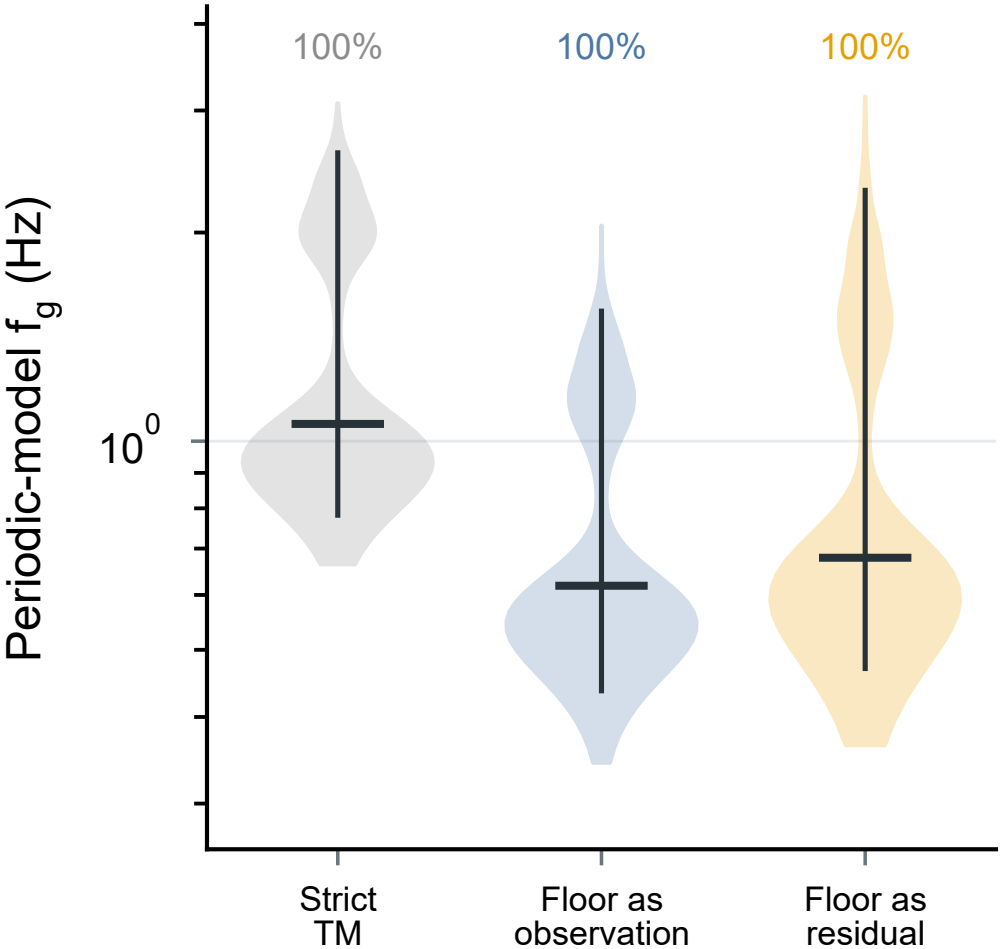
Biological uncertainty in model-derived gain turnover. Periodic-model *f*_*g*_ after jointly sampling functional branch weights and dynamical refits for the strict sensitivity control and the two residual-response interpretations. Violins show finite thresholds, bars mark medians and 95% intervals, and percentages give the fraction retaining turnover, rounded to the nearest whole percent. Each interpretation contains 5,000 joint draws; crossings were evaluated over 0.25–100 Hz.

### Joint waveform and polarity reliability at finite release-site counts

We evaluated the prespecified criterion of filtered NRMSE *<* 0.10 and polarity disagreement *<* 5% over 100 biological bootstrap parameter sets at 5, 10, and 20 Hz. The fraction of parameter– frequency combinations meeting both criteria increased from 32% at *N* = 20 to 57% at *N* = 30, 75% at *N* = 40, and 100% at *N* = 50 (Supplementary Fig. S7). These fractions summarize the finite-site reliability of the predicted output under the tested conditions.

**Figure S7:**
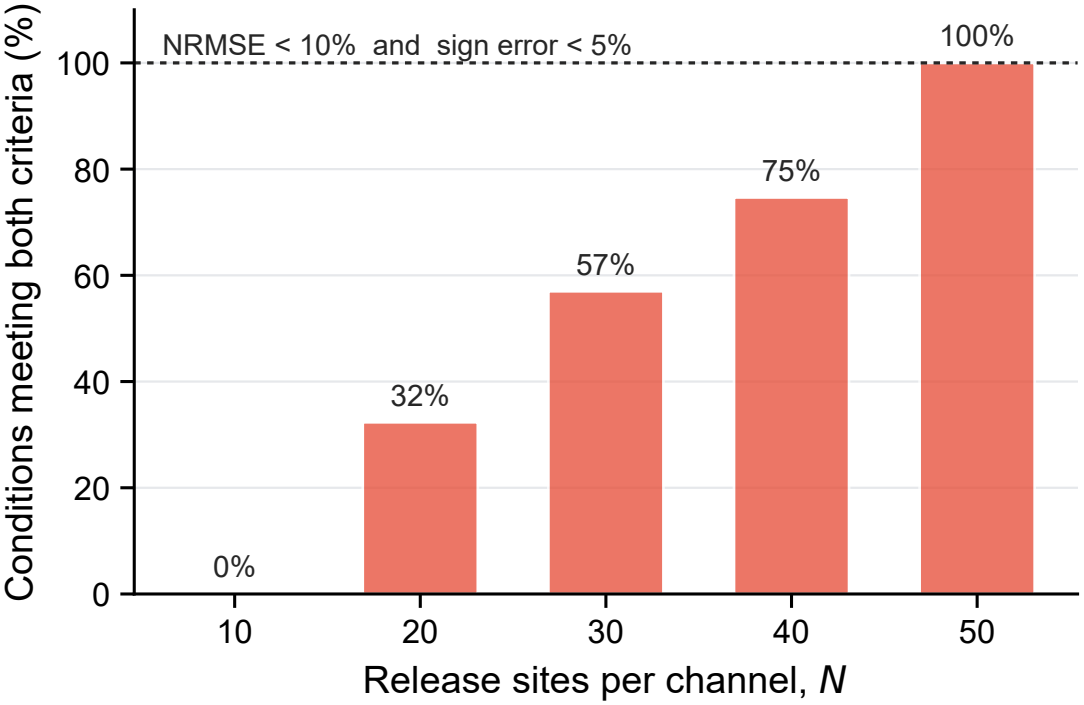
Joint waveform and polarity reliability improves with release-site number. Fraction of parameter–frequency combinations satisfying both filtered NRMSE *<* 10% and polarity disagreement *<* 5%, using the same 100 biological bootstrap parameter sets and three frequencies as Fig. 3c. Each bar summarizes 300 combinations; percentages are rounded to the nearest whole percent. All tested combinations meet the criterion at *N* = 50.

### Task-dependent effects of the co-release fraction

The SRNN co-release-fraction sweep showed task-dependent performance (Supplementary Fig. S8a–h). The best tested *α* ranged from 0.1 to 1.0 across datasets, and maximum gains ranged from 0.32 to 17.48 percentage points. The fixed value *α* = 0.5 provides a common comparison point across tasks. These task experiments complement the mechanistic results by showing that the same dual-branch computation can be exploited by learning, while leaving its optimal strength problem dependent.

**Figure S8:**
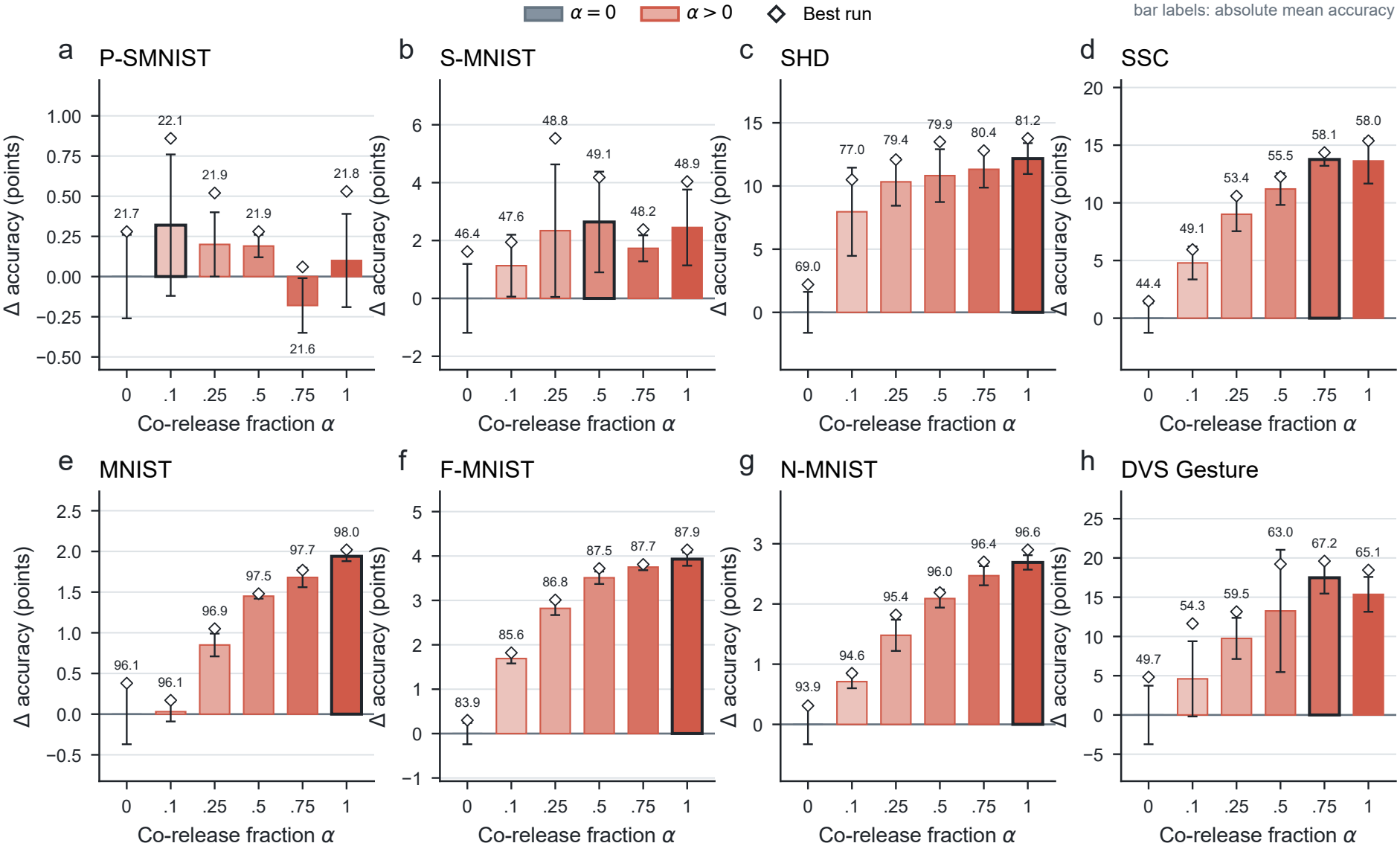
The co-release fraction associated with highest accuracy varies across tasks. (a– h) P-SMNIST, S-MNIST, SHD, SSC, MNIST, Fashion-MNIST, N-MNIST, and DVS Gesture, respectively. All six tested *α* values are shown separately for every dataset. Bars are supplied mean accuracies centered on the corresponding *α* = 0 mean, whiskers denote one supplied standard deviation of the raw accuracies, diamonds denote the supplied best run after the same centering, and numeric labels give absolute mean accuracy. The black-outlined bar marks the highest supplied mean among the tested values. Conditions at *α* = 0 and 0.5 contain *n* = 4 seeds; all other conditions contain *n* = 5. The highlighted *α* values identify the highest supplied mean within each tested grid.

## Supplementary Theory

### Local stability of the rate–resource system

At a fixed point of Eqs. 14–15, define

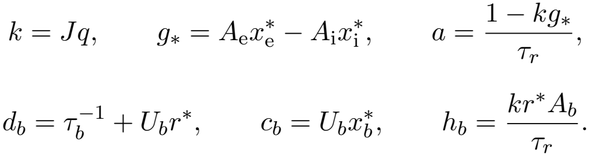

The Jacobian in the state order (*r, x*_e_, *x*_i_) is

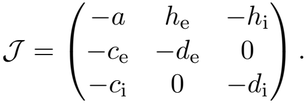

Expanding det(*sI* − *J*) gives *P* (*s*) = *s*^3^ + *a*_1_*s*^2^ + *a*_2_*s* + *a*_3_, with

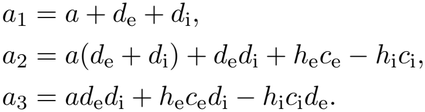

The Routh–Hurwitz criterion gives *a*_1_ *>* 0, *a*_2_ *>* 0, *a*_3_ *>* 0, and *a*_1_*a*_2_ *> a*_3_ as necessary and sufficient conditions for all Jacobian eigenvalues to have negative real parts. Under the local continuous-differentiability assumption, these conditions establish local exponential stability of the nonlinear system.

The equilibrium slope can be written as

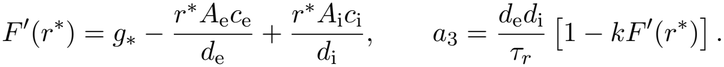

Negative equilibrium differential gain makes the constant term positive; the remaining inequalities determine the decay of all dynamical modes. The static-GABA implementation used in the reduced-model simulations has *U*_i_ = 0 and *x*_i_ = 1. On this invariant subspace the dynamics reduce to (*r, x*_e_); the three-state extension adds a stable inhibitory-resource mode with eigenvalue −1*/τ*_i_.

For Fig. 4a, equilibrium susceptibility was obtained from the working-point Jacobians as *χ* = −[*J* ^−1^***v***]_*r*_, where ***v*** = (*q/τ*_*r*_, 0, …)^⊤^. For each architecture, *F* denotes its corresponding equilibrium drive: *F*_static_(*r*) = (*A*_e_ − *A*_i_)*r, F*_common_(*r*) = (*A*_e_ − *A*_i_)*r/*(1 + *U*_e_*τ*_e_*r*), or the separate-resource expression in Proposition 4.5. Across all 273 model–working-point combinations at *J* = 8, the Jacobian response agreed with Eq. 16, using the corresponding *F* ^′^, to a maximum absolute difference below 10^−13^. The plotted normalized susceptibility includes stable fixed points only.

